# ABCF Protein-Mediated Resistance Shapes Bacterial Responses to antibiotics Based on their Type and Concentration

**DOI:** 10.1101/2025.02.10.637581

**Authors:** Markéta Koběrská, Ludmila Veselá, Michaela Novotná, Durga Mahor, Aninda Mazumdar, Nikola Pinďáková, Pamela Omena Petravicius, Julie Pokorná, Zdeněk Kameník, Gabriela Balíková Novotná

**Author notes:** authors contributed to the manuscript equally (Markéta Koběrská, Ludmila Veselá).

## Abstract

ABCF ATPases are increasingly recognized as translation factors that rescue stalled ribosomes, whether they encounter challenging mRNA templates or antibiotic-induced stalling. The latter defines ARE ABCF proteins, known for their role in antibiotic resistance. However, in this study, we reveal a broader role of ARE ABCFs in antibiotic-responsive regulation. Using genetic, OMICs, and biochemical approaches we showed that ARE ABCF proteins TiaA and Are5sc in *Streptomyces coelicolor* use their resistance functions to modulate specialized metabolism and proteosynthesis in response to lincosamide, streptogramin A, and pleuromutilin (LS_A_P) antibiotics. Although under LS_A_P exposure, either Are5sc or TiaA is essential for activating the biosynthesis of the redox-active antimicrobial actinorhodin, these proteins exhibit distinct functions at the proteome level, defined by their resistance profiles and temporally regulated expression. Are5sc facilitates early adaptive responses by modulating the WblC regulon across a broad range of LS_A_P concentrations, while TiaA is induced later, specifically at higher concentrations, where it suppresses antibiotic stress responses, particularly against pleuromutilins. TiaA function thus reflects the ecological context of LS_A_P antibiotics as pleuromutilins are produced by fungi, whereas lincosamides/streptogramins originate from actinomycetes. Our findings demonstrate that ARE ABCF proteins, through their resistance function, act as global regulators of translation, mirroring the roles of non-ARE ABCF proteins like EttA. This highlights their broader ecological and physiological significance, extending beyond their established role in antibiotic resistance.

**IMPORTANCE:** Bacteria adapt to diverse stimuli mainly through transcriptional changes that regulate adaptive protein factors. Here, we show that responses to protein synthesis-inhibiting antibiotics are fine-tuned by antibiotic resistance ABCF proteins at the translational level, enabling bacteria to differentiate between antibiotic classes and concentrations for a tailored response. Additionally, we have demonstrated that these proteins can specialize in conferring high-level resistance to specific antibiotics. Given their prevalence in pathogenic bacteria, ARE ABCF proteins may play a crucial role in resistance development, particularly against new antibiotics targeting the ribosomal catalytic center, presenting a significant challenge for antimicrobial therapy.

## INTRODUCTION

ABCF proteins are members of the ATP-binding cassette (ABC) family, a diverse group of ATPases involved in a broad range of cellular processes. Unlike canonical ABC transporters, which act as membrane-bound pumps, ABCF proteins operate in the cytoplasm and are increasingly recognized as key translational factors (1–9). These proteins rescue ribosomes stalled by challenging mRNA sequences, such as consecutive stretches of charged or bulky amino acid residues, as well as by ribosome-targeting antibiotics. ABCF proteins are widely distributed across bacteria, with multiple paralogs often encoded within the same genome, underscoring their functional importance (9). However, their broader physiological roles remain poorly understood.

A specific subset of ABCF proteins, known as antibiotic resistance ABCF (ARE ABCF) proteins, confers resistance to ribosome-targeting antibiotics by displacing them from the peptidyl transferase center (6, 7, 10, 11). These proteins are divided into subfamilies, each specialized in recognizing antibiotics that bind to a specific site on the ribosome (8, 9). Four of these subfamilies, including ARE5 subfamily, specifically respond to lincosamide streptogramin A, and pleuromutilin (LS_A_P) antibiotics that inhibit early steps of protein synthesis by binding to the peptidyl transferase center of the ribosome. While some ARE ABCF proteins regulate their own expression in response to targeted antibiotics, it is assumed that their primary function is generally associated with resistance to these antibiotics (5–14). However, our previous work has shown that the protein LmrC from the ARE5 subfamily, which is encoded in the lincomycin biosynthetic gene cluster (BGC) of *Streptomyces lincolnensis,* activates transcription of the *lmbU* gene in response to the same lincosamide-class antibiotic, thereby triggering premature lincomycin production (8). This finding reveals that the function of ARE ABCF proteins extends beyond antibiotic resistance to include antibiotic-responsive regulation that activates the biosynthesis of antibiotic encoded within the same BGC where LmrC is encoded. Specifically, ARE5 proteins are highly prevalent within the *Streptomyces* genus, with 95% of genomes containing at least one *are5* gene, and over a third containing multiple paralogs (9). While several ARE5 proteins are encoded within BGCs of ribosome-targeting antibiotics, indicating functional similarities to LmrC (8), the majority of ARE5 proteins are not linked to BGCs, presenting a significant challenge in predicting their physiological roles. In *Streptomyces coelicolor,* a model actinomycete with rich specialized metabolism and almost three dozen characterized BGCs, two ARE5 proteins that we named TiaA and Are5sc are encoded outside known BGCs.

This study shows, that TiaA confers resistance to inhibitory concentrations of tiamulin, suggesting that this mechanism protects the strain from pleuromutilin, a natural tiamulin analog produced by higher fungi in their shared soil environment. Moreover, Are5sc also confers resistance to LS_A_P antibiotics, but this function appears biologically redundant as TiaA can substitute Are5sc in all cases. However, we demonstrate that both Are5sc and TiaA utilize their resistance function to shape concentration dependent antibiotic response to different LS_A_P antibiotics. While both proteins are essential for inducing actinorhodin biosynthesis, each of these proteins plays distinct roles in shaping the overall antibiotic response, potentially reflecting different scenarios in the soil. Our study expands the limited research on the regulatory functions of ARE ABCF proteins, providing the first example of ARE5 proteins encoded outside BGCs shaping the antibiotic response. This finding, supported by proteomic and metabolomics data, suggests that antibiotic-responsive regulation could be a general function of these ubiquitous proteins, contributing to bacterial adaptation to their natural environments.

## RESULTS

### TiaA is essential for antibiotic resistance and strain protection, while Are5sc is indispensable

To elucidate the functions of the two ARE5 paralogs in *Streptomyces coelicolor* M145 (WT), we examined the antibiotic susceptibility of spores and mycelia of *tiaA* and *are5sc* single and double deletion mutants as well as mutants complemented with the respective genes (Fig. 1a and Supplementary Tables 1 and 2). A range of antibiotics targeting the large subunit of the bacterial ribosome, known to be influenced by ARE ABCF proteins, was tested. As expected, neither of the ARE5 proteins affected resistance to erythromycin, chloramphenicol and tetracycline – antibiotics that target ribosomal sites distinct from those targeted by LS_A_P antibiotics. However, the ARE5 proteins did influence susceptibility to LS_A_P antibiotics, with each paralog showing distinct effects. Whereas TiaA consistently provided high resistance to tiamulin (pleuromutilin) and pristinamycin IIA (streptogramin A). Its role in mediating resistance to lincosamides was evaluated using clindamycin (a semisynthetic lincosamide), as *S. coelicolor* is intrinsically resistant to lincomycin mediated by proteins of WblC regulon through WblC-regulated transcription of 312 genes, including *are5sc* (15). Resistance to clindamycin varied depending on whether *tiaA* was expressed under the native or constitutive promoter or co-expressed with *are5sc* (Fig. 1a). In contrast, Are5sc confers moderate resistance to clindamycin and pristinamycin IIA, but this effect was only evident in the absence of *tiaA*, suggesting a secondary or supporting role. We further confirmed the contribution of TiaA to tiamulin resistance by expressing *tiaA* in the *S. lincolnensis lmrC* deletion mutant, a strain naturally resistant to lincosamides and streptogramins A but susceptible to tiamulin (Supplementary Table 2). Our results demonstrate that TiaA is specialized for pleuromutilin resistance, while both TiaA and Are5sc contribute to resistance to lincosamides and streptogramins A. However, TiaA appears to be the dominant factor in conferring resistance to these LS_A_P antibiotics, as Are5sc is dispensable for resistance in the presence of TiaA.

**Figure 1.**
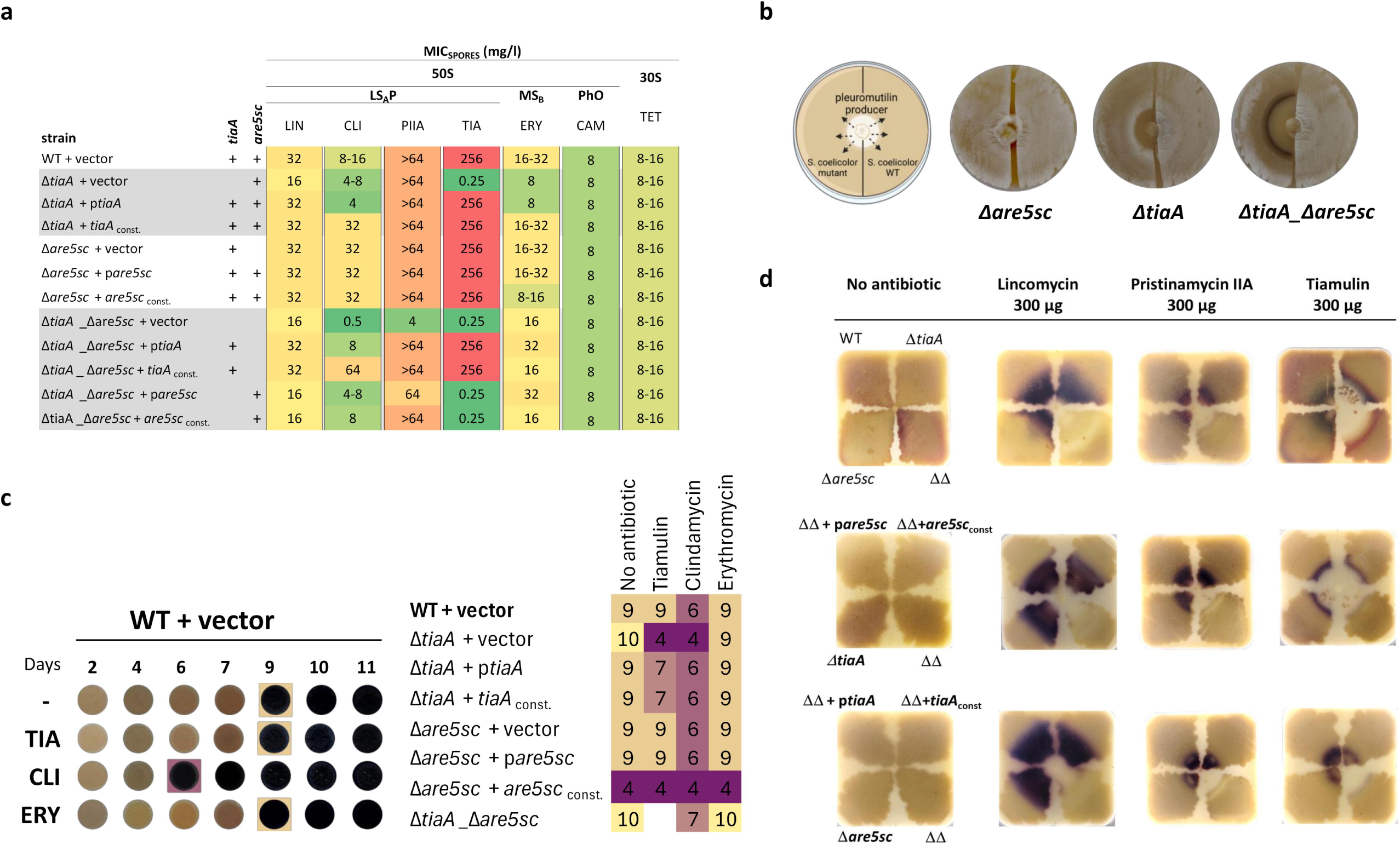
Phenotypic characterization of *tiaA* and *are5sc* mutants. **(a)** Heat map showing susceptibility of spores of WT, ARE5 deficient strains and complemented strainsto lincomycin (LIN), clindamycin (CLI), tiamulin (TIA), pristinamycin IIA (PIIA), erythromycin (ERY), chloramphenicol (CAM) and tetracycline (TET). The antibiotics are grouped based on resistance phenotypes conferred by ARE ABCF proteins: LS_A_P (lincosamides, streptogramin A, pleuromutilins), MS_B_ (macrolides, streptogramin B), and PhO (phenicols, oxazolidinones). **(b)** Co-culture of the pleuromutilin-producing basidiomycete C. passeckerianus CCBAS738 with S. coelicolor WT and *tiaA* and *are5sc* single and double mutants on MS agar (see also Supplementary Fig. 3) **(c)** The heatmap shows a day of the onset of actinorhodin production in response to sub-inhibitory concentrations (0.03 mg l^−1^ ) of TIA, CLI and ERY on MH agar in 6-well plates (WT shown as an example, full figure in Supplementary Fig. 2). TiaA modulates the timing of actinorhodin production, particularly in response to CLI and TIA. Constitutive overexpression of Are5sc results in premature actinorhodin production, independent of antibiotic exposure. Actinorhodin production was absent or weak in *tiaA_are5sc* mutant in the presence of TIA. **(d)** Production of actinorhodin around antibiotic discs with 300 µg of LIN or TIA shows that each antibiotic induces actinorhodin production in different concentration range depending on the strain susceptibility and that actinorhodin production in response to antibiotics is dependent on TiaA or Are5sc.

We hypothesize that TiaA’s specialization for resistance to tiamulin, a semisynthetic derivative of pleuromutilin produced by fungi of the genus *Clitopilus,* allows *S. coelicolor* to thrive in environments shared with these fungi. To test this hypothesis, *S. coelicolor* WT and mutant strains were co-cultured with two strains of *C. passeckerianus* – one that produces pleuromutilin and one that does not (Fig. 1b and Supplementary Fig. 1a). While both *C. passeckerianus* strains produced antibacterial compounds active against the *S. coelicolor* WT strain (Fig. S1a), only the pleuromutilin-producing strain exhibited an inhibition specifically associated with the absence of TiaA (Fig. 1b and Supplementary Fig. 1a). This inhibition indicates that TiaA plays a critical role protection from pleuromutilin, allowing *S. coelicolor* to survive and possibly compete with the pleuromutilin-producing fungus.

### Actinorhodin production in response to antibiotics is regulated by ARE5 proteins

In contrast to TiaA, which directly contributes to antibiotic protection, Are5sc is dispensable for this role. Instead, the resistance function of Are5sc suggests an antibiotic-responsive regulatory role, similar to LmrC. However, unlike LmrC, Are5sc is not associated with a BGC, leaving regulatory targets of Are5sc unclear. We initially focused on actinorhodin production, a blue pigment with redox-active properties(16), which is induced by ribosome-targeting antibiotics(17) through an unknown mechanism(18). Notably, actinorhodin production in response to pleuromutilin-producing fungi was negatively affected by TiaA (Supplementary Fig. 1b, c). To investigate further, we monitored the onset of actinorhodin production in WT and several deletion mutants and complemented strains of *S. coelicolor*, exposed to sub-inhibitory concentrations of LS_A_P antibiotics and erythromycin (Fig. 1c Supplementary Fig. 2). Consistent with previous reports(18), clindamycin induced earlier actinorhodin production in WT strain. TiaA deficiency not only increased sensitivity but also accelerated actinorhodin production in response to clindamycin and tiamulin (Fig. 1c and Supplementary Fig. 2). Are5sc deficiency, which does not affect antibiotic sensitivity, had no impact on the timing of actinorhodin production. However, overproduction of Are5sc accelerated actinorhodin production, even in the absence of an antibiotic. The absence of both ARE5 proteins resulted in the highest sensitivity to clindamycin and tiamulin and to impaired actinorhodin production in response to tiamulin. However, clindamycin-induced actinorhodin production was preserved with timing similar to the WT (Fig. 1c and Supplementary Fig. 2). These results demonstrate that TiaA delays the antibiotic-induced onset of actinorhodin production through its resistance function, while Are5sc appears to have a more complex regulatory role.

To determine the range of antibiotic concentrations that induce actinorhodin production, we used antibiotic discs to create concentration gradients on MS agar, a medium that supports complete differentiation, including sporulation, due to the use of mannitol as a poorly utilized carbon source (19) (Fig. 1d and Supplementary Fig. 3). Our results demonstrate that only the simultaneous deficiency of both ARE5 proteins severely impacted or completely abolished actinorhodin production in response to LS_A_P antibiotics, indicating a degree of functional redundancy between the two proteins in regulating this process. However, the specific antibiotic concentration range required to induce actinorhodin production varied depending on the ARE5 protein and the associated resistance it confers. Lincomycin and pristinamycin IIA induced actinorhodin production at similar concentration range regardless of whether TiaA, Are5sc, or both were present. In contrast, tiamulin induced actinorhodin production at higher concentrations in the presence of TiaA, whereas in the presence of only Are5sc, tiamulin triggered actinorhodin production within a narrow range of lower concentrations, specifically at the edge of the inhibition zone (Fig. 1d and Supplementary Fig. 3).

### ARE5 proteins fine-tune their expression that is induced by LS_A_P antibiotics through ribosome-mediated attenuation and that correlates with actinorhodin production

To gain insight into the relationship between LS_A_P antibiotic-induced actinorhodin production and ARE5 protein function, we investigated how expression of TiaA and Are5sc is regulated. Expression of ARE ABCF genes is often regulated by ribosome-mediated attenuation in response to antibiotics at the transcriptional level, as observed in *lmrC*, *vgaA*, *msrD*, *vmlR* and *vgaL* (5, 6, 8, 13, 20)This mechanism uses a small open reading frame (uORF) and a premature transcription terminator encoded within the 5’ untranslated regions (5’UTR). When ribosome-binding antibiotics inhibit translation of this uORF, a conformational switch from the terminator to the anti-terminator occurs, allowing full-length transcription of the gene(21). The large 5’UTR of *tiaA* (194 bp) and *are5sc* (232 bp) identified previously(22, 23) can indeed adopt terminator and anti-terminator conformations (Supplementary Fig. 4a, b). Additionally, analysis of the 5’UTRs of both genes using uORF4U(24) identified conserved overlapping uORFs in *are5sc* (uORFs *are5scL1* and *are5scL2*, encoding MLV and MRS leader peptides, respectively) and a single uORF in *tiaA* (*tiaAL*, encoding MVGDDDISG) (Fig. 2a). The ribosome binding sites of these uORFs overlap with anti-terminator hairpins (Supplementary Fig. 4a, b). Ribosome toeprinting experiments showed that only *are5scL1* and *tiaAL* are translated and inhibited by the LS_A_P antibiotics lincomycin, clindamycin, tiamulin and pristinamycin IIA but not by erythromycin (Fig. 2b, c and Supplementary Fig. 4c, d). Stalling at the start codon of these uORFs is consistent with the mode of action of LS_A_P antibiotics, which inhibit translation during early elongation steps. These results show that *are5scL1* and *tiaAL* are uORFs that are potentially involved in the attenuation-based regulation of *are5sc* and *tiaA*.

**Figure 2.**
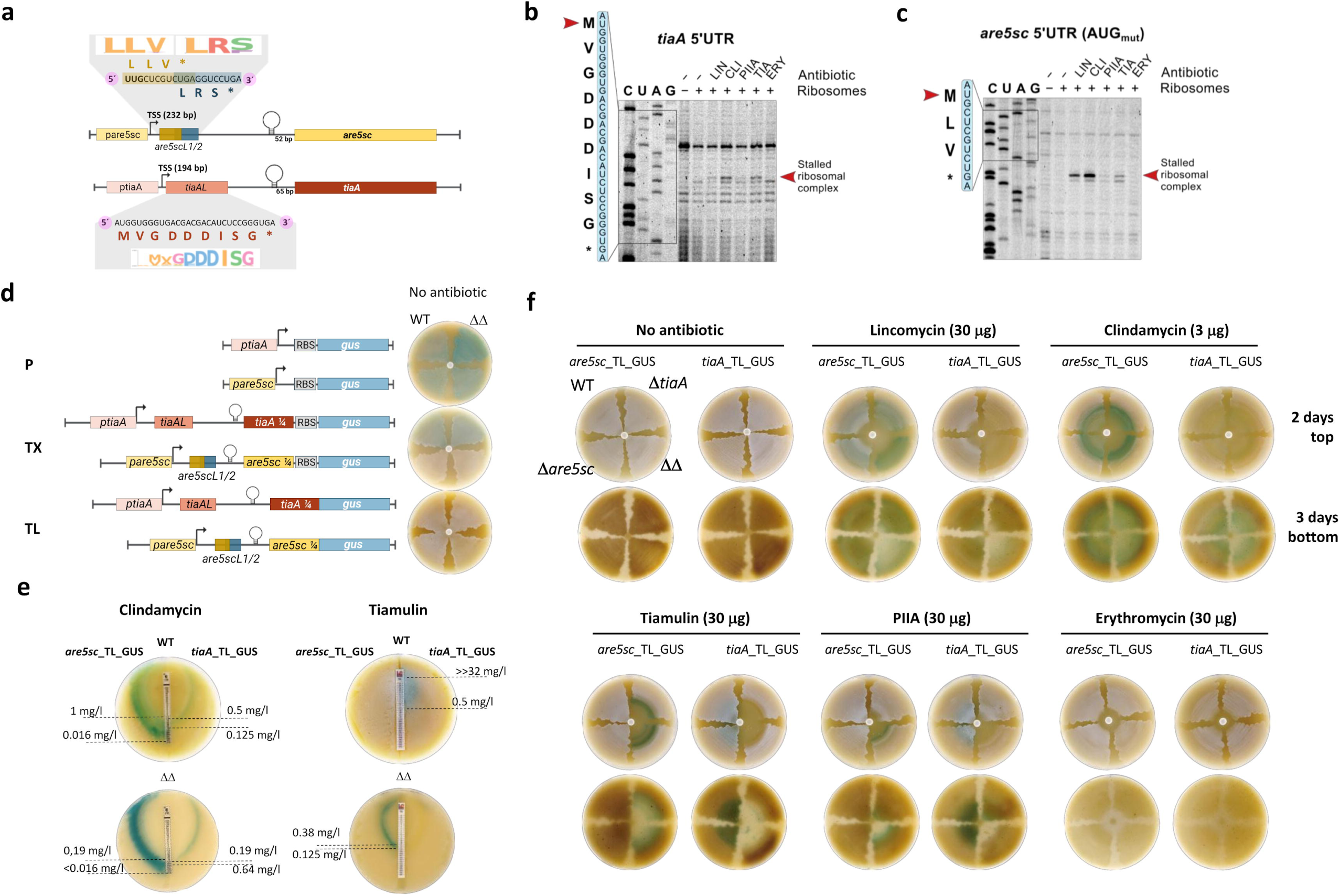
**TiaA and ARE5sc are expressed differentially in response to LS**_A_**P antibiotics (a)** Predicted small regulatory open reading frames (uORF) in the 5’ untranslated regions (5’UTR) of *are5sc* and *tiaA* genes and respective amino sequence logo showing conservancy across homologous ARE5 proteins. **(b,c)** Toeprint analysis using *Streptomyces venezuleae* ribosomes and an *E. coli* -derived *in vitro* reconstituted translation system (NEB PurExpress) performed on **(b)** *tiaA* 5’UTR **(c)** *are5sc* 5’UTR (AUG_mut_) mRNA templates in the presence of the LS_A_P antibiotics lincomycin (LIN, 50 µM), clindamycin (CLI, 5 µM), tiamulin (TIA, 50 µM), pristinamycin IIA (PIIA, 30 µM) and erythromycin (ERY, 50 µM). Toeprint bands indicating the position of stalled ribosome complex (SRC) are highlighted by red arrows. Toeprints with non-mutated *are5sc* template are shown in Suplementary Fig. 4c, d. **(d)** Promoter (P), transcriptional (TX), and translational (TL) reporter fusions were constructed to investigate the regulation of *tiaA* and *are5sc*. A β-glucuronidase (GUS) reporter activity in the absence of antibiotics indicates that *are5sc* is regulated at the transcriptional level, while *tiaA* seems to be regulated at the translational level (for explanation see Supplementary Fig. 5). **(e)** E-test strips were used to determine the range of antibiotic concentrations that induce the expression of *tiaA* and *are5sc* on MS agar **(f)** TL fusion reporter were used to monitor the expression of *tiaA* and *are5sc* in response to antibiotics diffusing from the paper discs placed in the center of MS agar plate inoculated by spores of WT, *tiaA* and *are5sc* mutants and double mutant (ΔΔ).

To determine whether ARE5sc regulation occurs at the transcriptional or translational level and to identify the antibiotics that induce its expression, we employed promoter (P), transcriptional (TX), and translational (TL) fusions with a β-glucuronidase (GUS) reporter (Fig. 2d). P-reporter constructs revealed that both promoters were constitutively active in the absence of antibiotics (Fig. 2d and Supplementary Fig. 5a). TX-reporters constructs showed that transcription of *are5sc* was induced by antibiotics, while *tiaA* was constitutive (Fig. 2d). The constitutive activity of the *tiaA* 5’ UTR TX-reporter was inconsistent with the predicted transcriptional attenuation mechanism. This discrepancy may be attributed to the use of a strong ribosome binding site in the reporter construct, which could potentially interfere with the native regulatory mechanism (Supplementary Fig. 5a, b). Only TL-reporters revealed that the production of TiaA and Are5sc is differentially triggered by antibiotics and that these proteins fine-tune their own expression (Fig. 2e, f). The expression of Are5sc was induced by lincomycin and clindamycin in all strains and by tiamulin and pristinamycin IIA only in the absence of TiaA, suggesting that TiaA negatively regulates the expression of Are5sc (Fig. 2f and Supplementary Fig. 5a). The induction of TiaA expression by lincosamides occurred one day later than that of Are5sc (Fig. 2f) and required a higher concentration (Fig. 2e). In contrast, induction by tiamulin and pristinamycin IIA occurred on the same day (Fig. 2f). However, the concentrations of tiamulin and pristinamycin IIA required to induce *tiaA* expression fall within the inhibition zone of the double mutant (Fig. 2e, f). Thus, *tiaA* expression relies on the resistance activity conferred by TiaA, or by Are5sc in the case of pristinamycin IIA.

Translational reporter assays also revealed a strong correlation between the presence of TiaA and Are5sc and actinorhodin production (Supplementary Fig. S6). The spatial overlap between GUS expression and antibiotic-induced actinorhodin production further supports the role of ARE5 proteins in regulating actinorhodin biosynthesis (compare Figs. 1d and 2f, Supplementary Fig. 6). All together, these findings highlight the complex interplay between ARE5 proteins, where their resistance functions not only influence their own expression but also impact the expression of other genes, as demonstrated by actinorhodin production in response to LS_A_P antibiotics.

### ARE5 proteins play a complex role in shaping the bacterial response to antibiotic presence

Even though ARE5 proteins are not encoded within the actinorhodin BGC, these proteins regulate actinorhodin biosynthesis in response to LS_A_P antibiotics, suggesting a broader role in antibiotic response that extends beyond ribosome protection. To investigate the global impact of ARE5 proteins on *S. coelicolor*, we performed a comprehensive proteomic analysis that compared the responses of WT, *tiaA* and *are5sc* deletion mutants, and overexpression strains to low and high subinhibitory concentrations of lincomycin, tiamulin, and erythromycin. Samples were collected at two time points following antibiotic induction, corresponding to early and late stages of specialized metabolite production (Fig. 3a). The WT strain exhibited distinct responses to different antibiotics and at different time points. Lincomycin triggered a delayed but robust response in which more than a thousand proteins were upregulated, including Are5sc and TiaA, as the most strongly induced proteins (Fig. 3b). Tiamulin, on the other hand, elicited a stronger immediate response at 24 hours, with Are5sc being the most significantly upregulated protein while TiaA was the most upregulated protein at 84 hours. Erythromycin, which does not induce ARE5 protein production, induced a similarly robust response as tiamulin at 24 hours but on the contrary to tiamulin minimum changes at 84 hours were detected (Fig. 3a, b). The WT did not show any significant changes in response to low antibiotic concentrations except Are5sc, which was the only significantly induced protein after lincomycin treatment (Supplementary Fig. 7a). The GUS reporter assay performed under the same cultivation conditions as proteomic analysis confirmed that ARE5 proteins production in response to antibiotics is concentration and time-dependent (Fig. 3c) which corresponds to similar observations on agar plates (Fig. 2f) and suggests that role of ARE5 proteins in antibiotic response is more complex.

**Figure 3.**
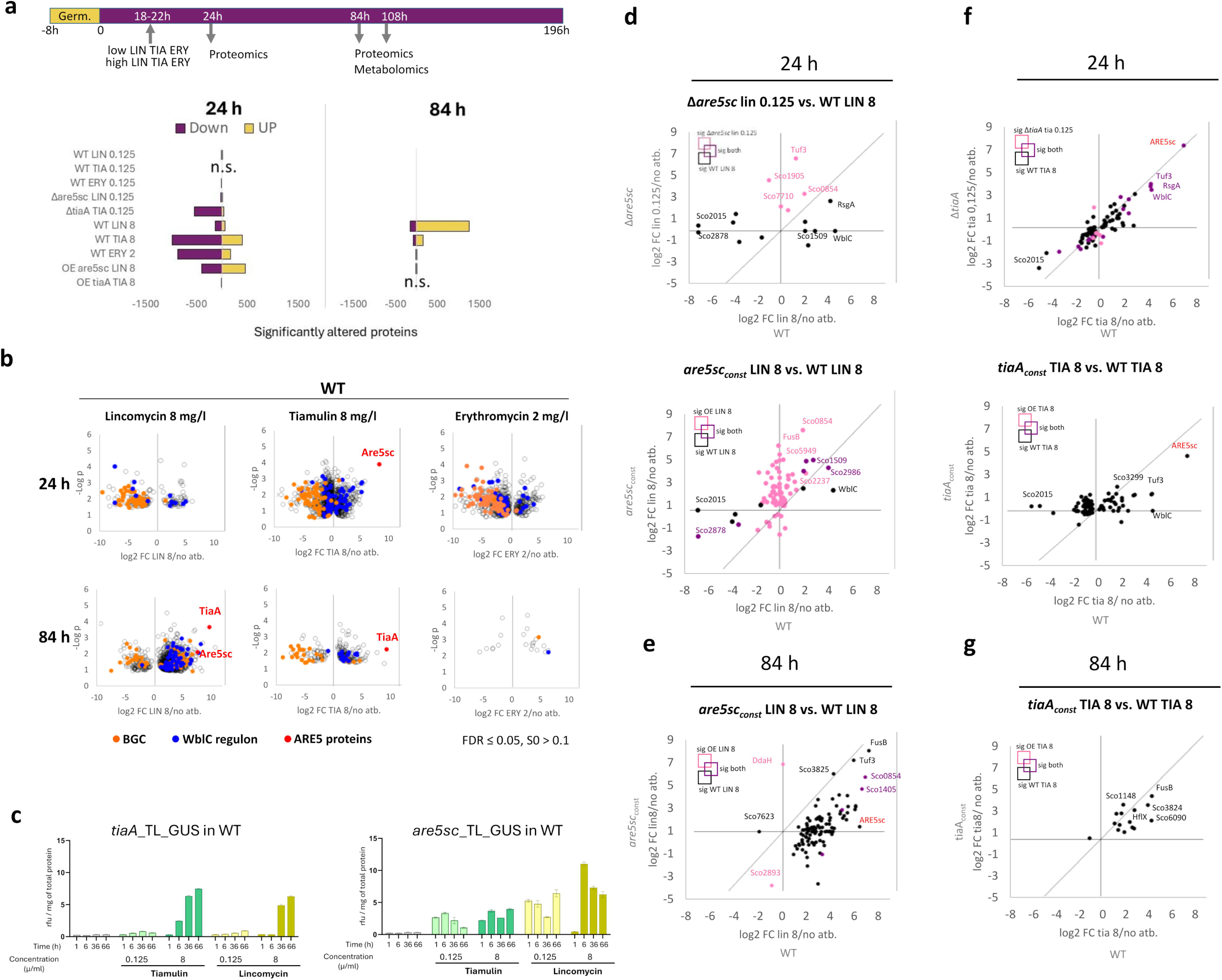
Proteomics analysis reveals the time-resolved production of TiaA and Are5sc and the effect of Are5sc on proteins of WblC regulon in *S. coelicolor* challenged by LS_A_P antibiotics. **(a)** *S*. *coelicolor* cultures were induced in the late exponential phase with low (0.125 mg/l) or high (8 mg/l) concentrations of lincomycin (LIN), tiamulin (TIA), or by erythromycin (ERY, 0.125 mg/l and 2 mg/l). Pellets and supernatants were collected at specific time points for analysis. The graph shows the number of significantly (FDR ≤ 0.05, S0 > 0.1) altered proteins for each analyzed pair of antibiotic-treated and untreated cultures. **(b)** Volcano plot showing significantly altered proteins in response to antibiotic treatment (FDR ≤ 0.05, S0 > 0.1). ARE5 proteins (red) are the most strongly induced proteins with Are5sc predominantly induced in 24 h and TiaA in 84 h. Production of proteins involved in specialized metabolite biosynthesis (orange) is delayed by high antibiotic concentration. **(c)** GUS reporter assays confirmed the time-dependent differential expression of ARE5 proteins under the same cultivation conditions as the proteomic analysis. **(d-g)** Scatterplots comparing the fold change in WblC regulon proteins abundance between antibiotic treated and untreated strains. The x-axis shows the fold change in one analysis and the y-axis shows the fold change in the other analysis. Proteins that are significantly changed in one (pink or black) or both (violet) analyzes are color-coded as indicated in each graph. Scatterplots comparing **(d)** WT and *Are5sc* mutant and overexpression (*are5*_const_ ) in 24h **(e)** in 84h **(f)** WT and *tiaA* mutant and overexpression (*tiaA*_const_) in 24h, **(g)** in 84.

Comparative analysis of significantly altered proteins in WT treated with lincomycin, tiamulin, and erythromycin revealed a conserved global response to antibiotics (Supplementary Fig. 8). Enrichment analysis revealed that this global response included downregulation of antibiotic biosynthesis and pyruvate metabolism, as well as the upregulation of the SigR regulon involved in the oxidative stress response(25, 26) (Supplementary Fig. 7b). On the other hand, significant differences were observed in proteins of the WblC regulon (Supplementary Figs. 7b and 8a). Tiamulin induced a stronger upregulation of WblC regulon proteins compared to lincomycin and erythromycin, with some proteins being uniquely upregulated by tiamulin and lincomycin (Supplementary Fig. 8).

Comparative analysis of significantly altered proteins of the WblC regulon in WT and the Are5sc deficient or overexpression strain revealed substantial changes driven by Are5sc (Fig. 3d-g). Loss of Are5sc resulted in the upregulation of specific WblC regulon proteins in response to low concentrations of lincomycin, suggesting that Are5sc negatively regulates their production. Overexpression of Are5sc led to the upregulation of a broader set of WblC regulon proteins, indicating the stimulatory effect of this protein (Fig. 3d). This effect was transient, as WblC regulon proteins were not upregulated in 84 hours which ruled out the possibility that upregulation of WblC regulon is due to a translational stress induced by Are5sc overexpression (Fig. 3e).

In contrast, TiaA functions mainly as a resistance protein and suppresses the antibiotic response. TiaA deficiency resulted in an enhanced response to low concentrations of tiamulin that mimicked the WT response to high concentrations (Fig. 3f). Conversely, overexpression of TiaA suppressed the response to tiamulin and resulted in minimal proteomic changes (Fig. 3f, g). These findings highlight the distinct roles of Are5sc and TiaA in modulating the bacterial response to LS_A_P antibiotics. While Are5sc appears to fine-tune the production of WblC regulon proteins, TiaA primarily functions as a resistance protein. The temporarily staggered production of ARE5 proteins allows for a coordinated response to antibiotic presence, with Are5sc initially adapting the cell to the antibiotic and TiaA subsequently neutralizing the antibiotic’s effect.

### TiaA mitigates the negative effect of LS_A_P antibiotics on quorum sensing-driven production of specialized metabolites

In our proteomic analysis, gene enrichment revealed a negative impact of antibiotic treatment on specialized metabolism at 24 hours, particularly disrupting the biosynthesis of coelimycin P1, calcium-dependent antibiotic (CDA), and undecylprodigiosin (Supplementary Fig. 7b). This finding suggests that, in response to antibiotics, ARE5 proteins may regulate the production of other specialized metabolites, in addition to actinorhodin. To investigate further, we performed metabolomic analysis on late-stage supernatants from the same cultures used for proteomics by liquid chromatography mass spectrometry (LC-MS) and correlated the identified metabolites (Supplementary Fig. 9b) with the abundance of known biosynthetic proteins. Proteins associated with biosynthetic pathways downregulated at 24h were detected at later time points (Fig. 4a) and their corresponding metabolites were detected at lower amounts than in WT. This downregulation is likely mediated by the inhibition of the gamma-butyrolactone (GBL) signaling pathway(27), as key components of this pathway were also suppressed following antibiotic treatment (Fig. 4b). Notably, Are5sc did not affect enzymes of the specialized metabolite biosynthetic pathways, as antibiotic-induced changes in protein abundance in the *are5sc* mutant and overexpression strain mirrored those observed in the wild-type strain (Fig. 4b). In contrast, TiaA played a distinct role in counteracting the negative effects of tiamulin on biosynthetic proteins. In the TiaA overexpression strain, the adverse impact of tiamulin on BGCs was neutralized, whereas absence of TiaA led to the downregulation of biosynthetic pathways, even at low tiamulin concentrations. These findings highlight a specific role for TiaA in mitigating the general antibiotic response to tiamulin, which includes delaying the onset of coelimycin P1, CDA, and prodiginine biosynthesis.

**Figure 4.**
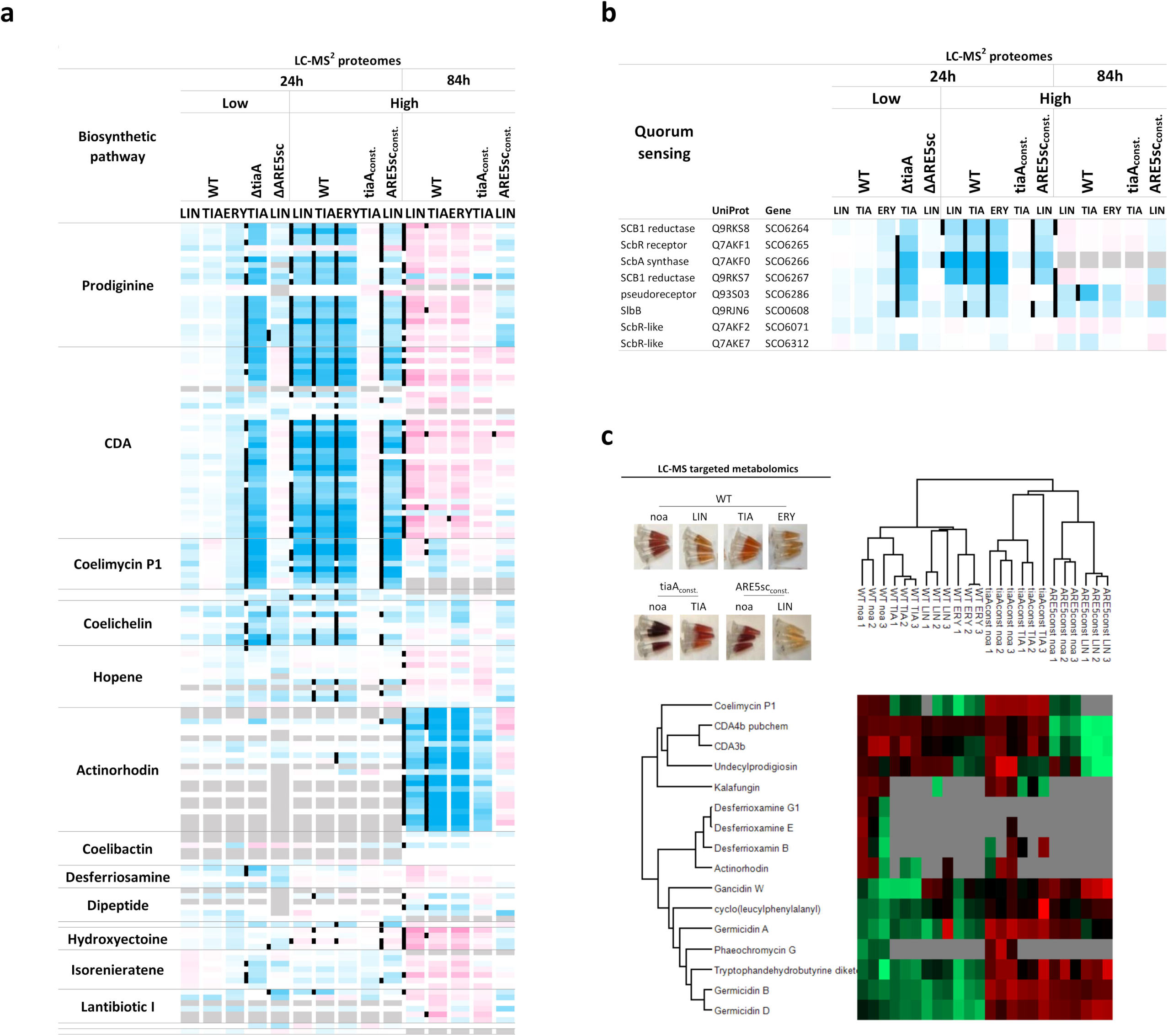
Proteomics and metabolomics analysis reveals the impact of TiaA on specialized metabolism in *S. coelicolor* challenged by LS_A_P antibiotics. **(a)** Heatmap illustrates the relative abundance ratios (log2) of proteins involved in known specialized metabolic pathways in *S*. *coelicolor* treated with antibiotics compared to control cultures. Proteins exhibiting significant differential abundance (FDR ≤ 0.05, S0 > 0.1) are highlighted with black squares. **(b)** Heatmap illustrates the relative abundance of proteins involved in GBL signalling pathways in response to antibiotic treatment. **(c)** Supernatants for LC-MS analysis of specialized metabolites were collected from the same cultures used for proteomic analysis, ensuring a direct correlation between protein expression and metabolite production. A clustering analysis of relative metabolite abundance (normalized peak intensities) in supernatants of wild-type and overexpression strains treated with high concentrations of antibiotics.

Actinorhodin is a late-stage secondary metabolite which, along with its intermediate kalafungin, was detected at high levels in the WT and TiaA-overexpressing strains without antibiotic treatment and at low level in TiaA-overexpresing strain or only sporadically in antibiotic-treated WT (Fig. 4c). While the Are5sc-overexpressing strain showed a slight and non-significant upregulation of actinorhodin biosynthetic proteins (Fig. 4a), actinorhodin was not detected in the supernatant (Fig. 4c). These findings suggest that neither the antibiotic concentrations used nor the overexpression of Are5sc induced actinorhodin production in *S. coelicolor* grown in YEME broth.

### Metabolic profile of WT and *tiaA*- and *are5sc*-overexpressing strains substantially differs

While proteomics primarily identified changes in pathways mentioned above, metabolic profiling revealed broader effects. Antibiotic treatment negatively affected the production of desferrioxamine in the WT strain and phaeochromycin G in both the WT and TiaA-overexpressing strains. (Fig. 4c and Supplementary Fig. 9). More importantly, lincomycin, and to a lesser extent tiamulin, stimulated the production of diketopiperazines, including gancidin W, cyclo(leucylphenylalanyl), and tryptophan dehydrobutyrin. (Supplementary Fig. 9b). This increased production correlates with the upregulation of leucine biosynthesis proteins in lincomycin-treated samples (Supplementary Fig. 7b). The production of cyclic peptides, and metabolites derived from acetate pathway phaeochromycin G and germicidines (28) were produced independently of the presence of antibiotic at higher levels in *tiaA*- and *are5sc*-overexpressing strains than in WT. These findings suggest that ARE5 proteins may indirectly influence the production of these specialized metabolites, potentially through metabolic reprogramming or altered cellular physiology.

### Pleuromutilin resistance evolved within multiple phylogenetic groups of ARE5 proteins

Phylogenetic analysis of the ARE5 protein subfamily *within Actinomycetota* revealed distinct taxonomic lineages, with at least five groups characteristic of *Streptomyces* genus including Are5sc and TiaA clades (Fig. 5a and Source data file). However, TiaA homologs are also often present across various *Actinomycetota* genera (Fig. 5b). This finding suggests that TiaA may have undergone horizontal gene transfer, which could explain its role in pleuromutilin resistance which is further supported by lower conservation of TiaA genomic context compared to the Are5sc lineage (Supplementary Fig. 10). To evaluate the prevalence of pleuromutilin resistance, we tested a collection of *Actinomycetota* strains with available genomic data, most of which harbor two ARE5 paralogs (Fig. 5b, Source data file). The tested strains exhibited higher levels of resistance to tiamulin compared to other antibiotics (Fig. 5b, c), suggesting an adaptive need to protect against this antibiotic class produced by fungi. Among seven strains containing TiaA homologs, three displayed high tiamulin resistance with MIC values of 256 mg/L. Interestingly, four strains with lower MIC values (16–64 mg/L) exhibited variations in the amino acid sequence within the extension of the antibiotic resistance domain (ARD) (Supplementary Fig. 11). Amino acid substitutions in this region are known to influence resistance levels in ARE ABCF proteins, as demonstrated for VgaA(29). Furthermore, seven additional strains lacking TiaA homologs but possessing other ARE5 proteins also showed elevated resistance to tiamulin (MIC 128–256 mg/L). This finding suggests that pleuromutilin resistance has independently evolved multiple times within distinct phylogenetic groups of the ARE5 subfamily (Fig. 5d).

**Figure 5.**
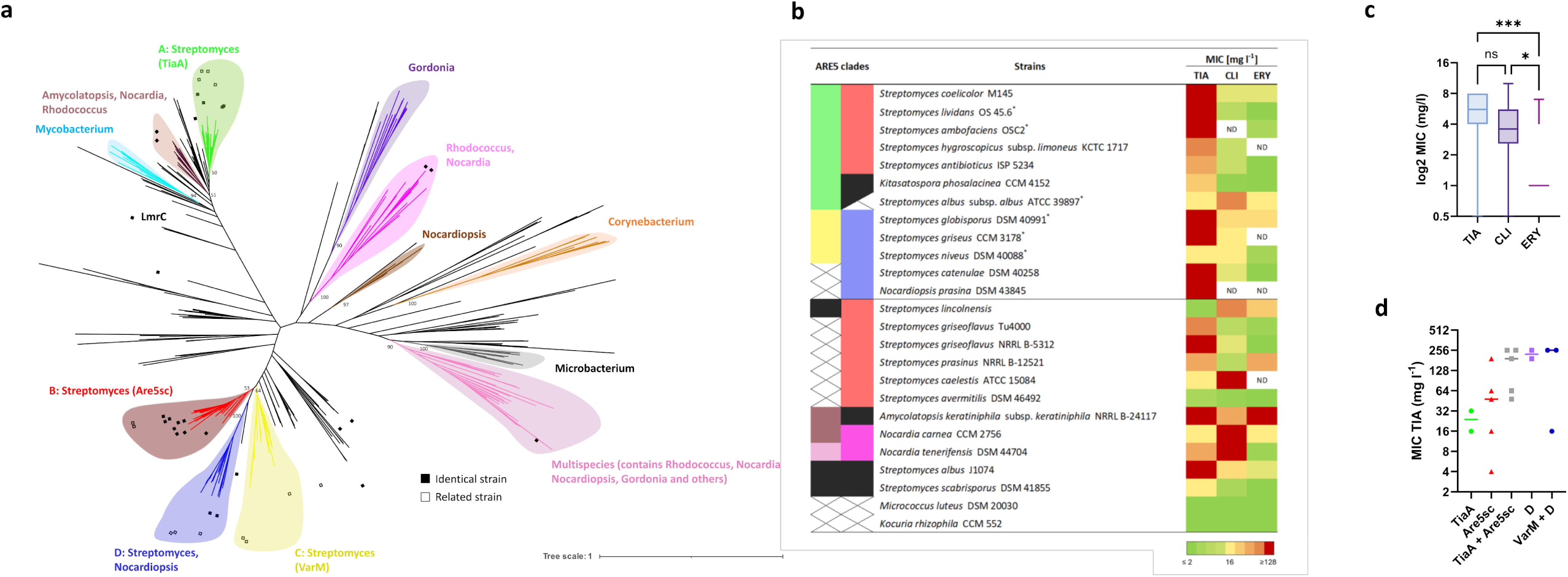
Specialized resistance to pleuromutilin evolved within different phylogenetic lineages of *Actinomycetota*. **(a)** A phylogenetic tree of ARE5 proteins as identified in (9). Different clades contain proteins from the same taxonomic groups. ABCF proteins from strains tested for antibiotic resistance are indicated by black squares, ABCF proteins from the same species but from different tested strain are indicated by open squares. **(b)** Heatmap showing the minimum inhibitory concentrations (MIC) of tiamulin (TIA), clindamycin (CLI) and erythromycin (ERY) for *Actinomycetota* strains. Most strains encode two ARE5 proteins from different clades, indicated by the same color as in (a). **(c)** Boxplots showing the distribution of MICs. **(d)** The distribution of MICs of tiamulin across clades shows that the highest MICs occur in strains encoding either TiaA in combination with Are5sc or ABCF proteins from clade D

## DISCUSSION

We have uncovered a complex interplay between Are5sc and TiaA proteins which shapes the bacterial response to LS_A_P antibiotics. Our findings reveal that ARE5 proteins play a broader regulatory role in antibiotic response, extending beyond their previously recognized association with BGC-encoded LmrC(8). While both Are5sc and TiaA are critical for the activation of actinorhodin production in response to LS_A_P antibiotics (Fig. 1d) they exhibit distinct roles at the proteome level (Fig. 3d-g). Are5sc, a weak resistance protein (Fig. 1a), is rapidly induced at low antibiotic concentrations (Figs. 2d-f and 3c) and modulates the WblC regulon (Fig. 3d). In contrast, TiaA, a specialized pleuromutilin resistance protein (Fig. 1a), is induced later and at higher antibiotic concentrations (Figs. 2d-f and 3c), where it suppresses the global antibiotic response (Fig. 4a and Supplementary Fig. 8d). This sequential and concentration-dependent induction of ARE5 proteins enables S. coelicolor to dynamically adapt to the LS_A_P antibiotics. Are5sc fine tunes the adaptive response at low concentrations, while TiaA neutralizes the antibiotic’s effect at higher concentrations. This behavior aligns with the general concentration-dependent effects of antibiotics on bacteria: low concentrations trigger adaptive responses, while high concentrations inhibit growth or induce cell death (17, 18, 30, 31). This phenomenon mirrors the natural occurrence of antibiotics in the environment, where diffusion from a producing organism creates a concentration gradient that might intensify over time (32). Here, we show that a pair of resistance proteins, Are5sc and TiaA, mediate this adaptive response in S. coelicolor. Furthermore, the pairwise occurrence of ARE5 proteins observed in about one third of bacterial genomes (8, 9) suggests that this paired system is a conserved and widespread strategy for recognition and adaptation to LS_A_P antibiotics in Actinomycetota.

Correlation between regulatory and resistance functions suggests that ARE5 proteins regulate gene expression through their abilities to confer resistance, i.e. by recognizing and displacing specific antibiotics from the ribosome (5, 6, 10–14, 29, 33). We have shown that TiaA and ARE5sc modulate each other’s expression through their resistance activities (Fig. 3d). This interplay results in distinct induction profiles for each ARE5 protein depending on the type and concentration of antibiotics (Fig. 6). Similarly other studies have shown that ARE ABCF proteins, such as VgaA and MsrD, can positively or negatively self-regulate their expression through a mechanism involving transcriptional attenuation, according to their antibiotic resistance function (5, 13). We hypothesize that the same regulatory principle leveraging the resistance function of ARE ABCF proteins is also used to modulate the expression of other genes, such as in the regulation of *lmbU* by LmrC (8). This suggests that ARE ABCF proteins represent a conserved mechanism for fine-tuning the bacterial response to antibiotic signals or stress in different microorganisms encoding these proteins (10, 11, 34).

**Figure 6:**
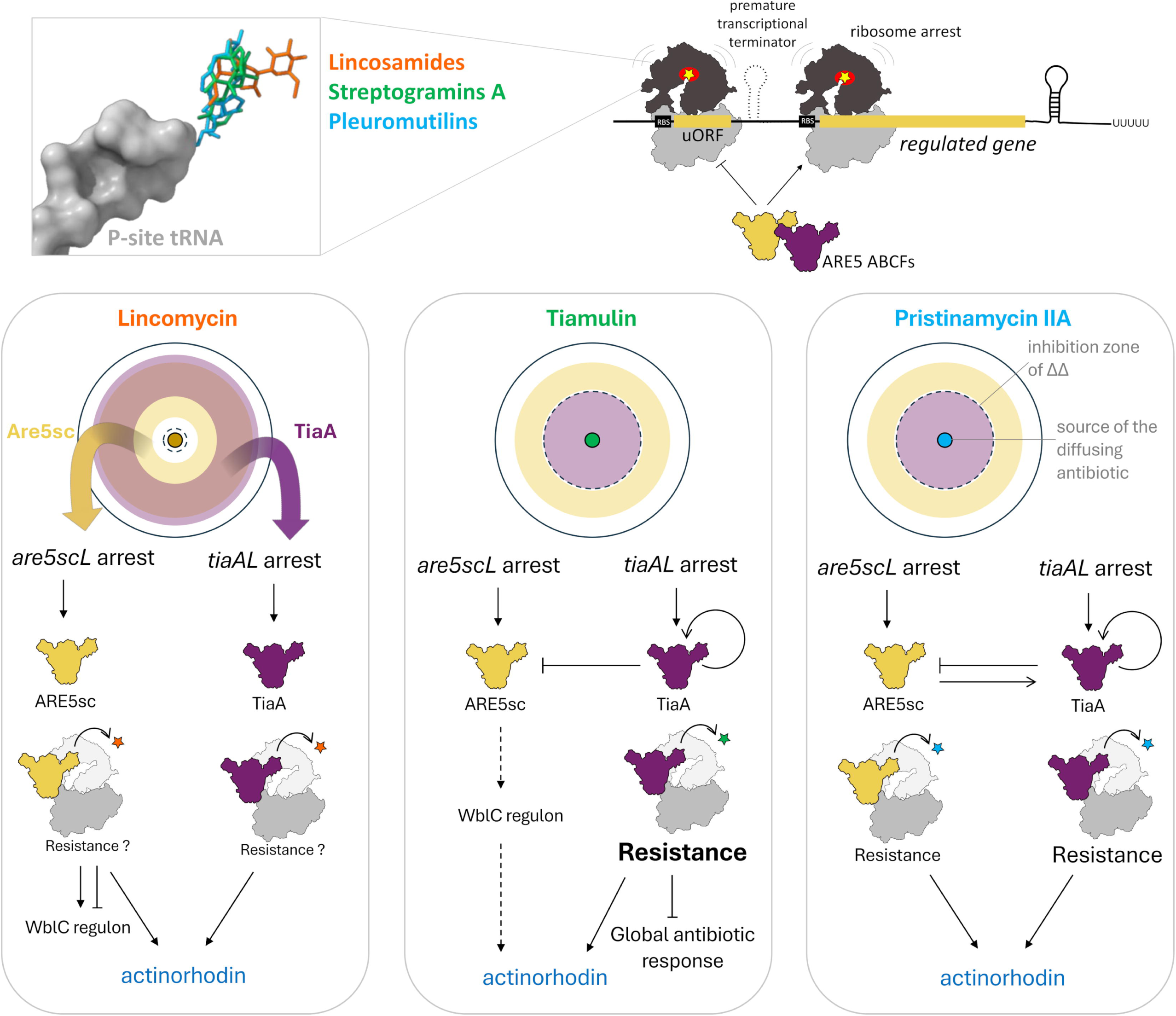
Interplay of TiaA and Are5sc in shaping the antibiotic response. A model illustrating the dynamic interplay between ARE5 proteins in modulating the bacterial response to LS_A_P antibiotics in a natural environment, where antibiotics diffuse from their source, creating a concentration gradient. LS_A_P antibiotics bind to the ribosomal peptidyl transferase center and stall ribosomes at the start codon. Ribosome stalling at upstream open reading frames (uORFs) induces anti-terminator conformations in the 5’ untranslated regions (5’ UTRs) of *are5* genes, initiating their transcription. The areas of ARE5 protein production on plates with a concentration gradient of antibiotics are represented in yellow for Are5sc and violet for TiaA. ARE5 proteins, guided by their resistance profiles, regulate their own expression and that of other genes, including those in the WblC regulon, via positive (arrows) and negative (blunt-ended lines) feedback loops. Their expression is induced by antibiotics in a type- and concentration-dependent manner. Are5sc, a weaker resistance protein, is activated at lower antibiotic concentrations and modulates the expression of WblC regulon proteins, enabling early adaptive responses. Conversely, TiaA, a stronger resistance protein, is induced at higher antibiotic concentrations, where it suppresses the global antibiotic response, particularly against pleuromutilins. As Are5sc does not confer significant resistance to tiamulin, its regulation of actinorhodin production likely involves indirect modulation of WblC regulon proteins (indicated by dashed arrows).

Recently, the ABCF protein EttA was shown to play a global regulatory role in *Escherichia coli* by modulating the translation of specific mRNAs encoding TCA enzymes (4). Similar to ARE ABCF proteins, EttA rescues stalled ribosomes. However, EttA specifically rescue stalling that is triggered by specific patterns of negatively charged amino acids within the nascent polypeptide chain rather than by the antibiotic binding (4). Nevertheless, both studies highlight the potential of ABCF proteins to globally modulate translation and thus provide an additional level of regulation that can significantly contribute to the organism’s adaptation to fluctuating external conditions.

The strength and specificity of resistance conferred by LS_A_P-responsive ARE-ABCF proteins can be fine-tuned through amino acid substitutions within the antibiotic-resistance ARD domain that reaches the antibiotic binding site (12, 29, 35). This demonstrates that the resistance function of ARE-ABCF proteins can evolve in response to selection pressure as evidenced by identification of at least two independent lineages of ARE5 proteins specialized for pleuromutilin resistance (Fig. 5). Understanding the molecular mechanisms leading to resistance specialization of ARE ABCF proteins is particularly critical, especially since these proteins are frequently present in clinically important microorganisms. Their specialization may be accelerated due to selection pressure from novel LS_A_P antibiotic derivatives which were recently developed or approved (36–38).

The ability of ARE5 proteins to discriminate between functionally similar LS_A_P antibiotics, which would otherwise have the same effect on cells, allows *S. coelicolor* to respond to these compounds based on their specific ecological context. Lincomycin, that is like other lincosamides produced by *Actinomycetota*(39, 40) elicits a stable, long-term response, including a strong induction of actinorhodin production (Figs. 1d and 3a). This response could be related to competition with closely related species, which is associated with the induction of antibiotic production (41). In contrast, fungal-derived pleuromutilin (42) elicited a stronger, more immediate response that was less sensitive to the induction of actinorhodin production, perhaps reflecting a defense strategy against a perceived threat from another domain of life (43, 44).

Our findings suggest that Are5sc modulates the expression of WblC regulon proteins in response to antibiotic stress. Unlike the SigR regulon, which exhibited a uniform response to the tested antibiotics (Supplementary Fig. 7b), the WblC regulon displayed a more nuanced, context-dependent response, with specific proteins being upregulated or downregulated based on the antibiotic and strain (Fig. 3d, e). Previously, the WblC regulon was defined by its transcriptional response to sub-inhibitory concentrations of tetracycline and the binding of WblC to 312 promoter regions (15). However, our proteomic analysis revealed that only a subset of these genes was upregulated in response to LS_A_P antibiotics, either as an early or late response (Fig. 3d, e and Supplementary Fig. 7b). This finding indicates that WblC regulon expression may be further modulated at the translational level depending on the specific type of stress. WblC itself is regulated *via* ribosome-mediated attenuation and responds to a broad spectrum of ribosome-targeting antibiotics and amino acid starvation (45). Notably, WblC protein expression was detected only at higher antibiotic concentrations in the wild-type strain and was absent in the Are5sc overexpression strain, suggesting that Are5sc negatively regulates WblC expression. Interestingly, while WblC was not upregulated in the Are5sc overexpression strain, numerous WblC regulon proteins were induced by lincomycin (Fig. 3d). This observation implies that Are5sc may regulate these proteins independently of WblC, potentially at the translational level. These findings highlight a complex regulatory network, wherein Are5sc fine-tune bacterial responses to antibiotic stress that is at transcriptional level regulated by WblC. Future studies will be essential to identify the direct targets of Are5sc and to uncover the molecular mechanisms by which WblC regulon proteins are selectively regulated under varying antibiotic conditions. Such insights could provide a deeper understanding of bacterial adaptation strategies and innate resistance mechanisms.

## MATERIALS AND METHODS

### Bacterial strains

The bacterial and fungal strains used in this study are listed in Supplementary Table 3. The WT strain was *S. coelicolor* M145 (46). *E. coli* XL10 (Agilent) and *E. coli* DH5α (Invitrogen) are used for routine cloning. E.⍰Zcoli BW25113/pIJ790 and *E. coli* DH5α/BT340 were used for generating deletion mutant *ΔtiaA*. *S. venezuleae* ATCC 10712 was used as a source of ribosomes for toeprinting experiments (47) E.⍰Zcoli ET12567/pUZ8002 was used for the intergenic conjugation from E.⍰Zcoli to S.⍰Zcoelicolor. Fungal strains *C. passeckerianus* CCBAS738 and CCBAS739 were acquired from Culture Collection of Basidiomycetes of the Institute of Microbiology of ASCR (https://www.biomed.cas.cz/ccbas/fungi.htm). *Actinomycetota* strains were obtained from the specified collections (Supplementary Table 3) and have been long-term preserved in our laboratory collection.

The in-frame deletion mutant *ΔtiaA* (SCO0636) was generated using the PCR-targeting method (48) by replacing the entire coding sequence of the *tiaA* gene in cosmid 7G01 with a cassette 773, which carries the apramycin resistance gene (*aac(3)IV*) and the *oriT* region of plasmid RK2, flanked by FRT sites. Primers Sco0636FW and Sco0636Rev used for disruption of the *tiaA* gene are listed in Supplementary Table 4. The mutated cosmid 7G01-*tiaA*::*apra* was introduced into *S. coelicolor* M145 by conjugation. The kanamycin-sensitive (Kan^S^), apramycin-resistant (Apra^R^) double-crossover mutant strain Δ*tiaA* with replaced gene by the *773* cassette was confirmed by PCR. I The in-frame deletion mutant Δare5sc (SCO6720) was generated using the CRISPR-Cas9 genome editing protocol for actinomycetes(49). The gene was deleted in such a way that encoding sequence for five amino acids from the N-terminus and five from the C-terminus were retained. The single-guide RNA (sgRNA) sequence was identified using the CRISPy software (50). The target sequence (*ATGCCCAGCGACTCCAGGAA* ) was amplified with primers sgRNA_R and sgRNA_6720_F, digested with *NcoI* and *SnaBI* restriction enzymes, and cloned into the pCRISPR-Cas9 vector. A homologous recombination (HR) template was constructed using two approximately 1 kb fragments flanking the 5’ and 3’ regions of the *are5sc* gene in plasmid pMK048 which was conjugated into *S. coelicolor* M145. Exconjugants were incubated with thiostrepton (1 mg/l) to induce the Cas9 enzyme under the control of the *tipA* promoter. The loss of the pCRISPR-Cas9 plasmid was facilitated by incubating the mutants at 37 °C for 24 hours to eliminate the temperature-sensitive vector. Colony PCR was used to screen for deletion mutants. Modified regions were further confirmed by Sanger sequencing. The double deletion mutant *ΔtiaA_Δare5sc* was generated by replacing *tiaA* with the inactivation cassette 773, using the mutated cosmid 7G01-*tiaA::apra* in the *Δare5sc* deletion mutant, as described above.

### Plasmid construction

All plasmids used in this study, including a description of the construction, are listed in Supplementary Table 5. For complementation of ARE5 mutants *in trans* we used integrative constructs with the respective genes placed either under the constitutive *ermEp* promoter (tiaA_const_ and are5sc_const_) or their native promoters (ptiaA and pare5sc). In tiaA_const_, the coding sequence of *tiaA* gene was cloned between the *NdeI* and *XhoI* sites of vector pIJ10257 (51), while in are5sc_const_, the *are5sc* gene coding sequence was inserted between the *NdeI* and *HindIII* sites of the same vector. In construct ptiaA and pare5sc, the *tiaA* gene, including its 400 bp upstream region and the *are5sc* gene, including its 323 bp upstream region, respectively, were inserted into *Eco*RV site of the promoterless vector pMS81 (52).

For promoter fusion (P) constructs, the strong synthetic ribosome-binding site (RBS) N2 and the *gus* gene encoding β-glucuronidase (53) were inserted into the *ptiaA* plasmid, retaining a 259 bp region of the *tiaA* promoter (plasmid *tiaA_P_GUS*), and into the *pare5sc* plasmid, retaining a 130 bp region of the *are5sc* promoter (plasmid *are5sc_p_GUS*). For transcription fusion (TX) constructs, RBS N2 and *gus* were inserted into the *ptiaA* and *pare5sc* plasmids, retaining the entire upstream regions of *tiaA* and *are5sc*, along with one-third of the respective coding sequences (plasmids *tiaA_TX_GUS* and *are5sc_TX_GUS*). For translation fusion (TL) constructs, the RBS N2 sequence in *tiaA_TX_GUS* and *are5sc_TX_GUS* plasmids was replaced in-frame with a sequence encoding the PGGGS linker, creating the constructs *tiaA_TL_GUS* and *are5sc_TL_GUS*. All plasmids were verified by Sanger sequencing and introduced into recipient strains via conjugation (54). Primers used for plasmid construction are listed in Supplementary Table 4.

### Growth media and cultivation conditions

Cultivation and sporulation of *Streptomyces* species was done according to standard protocols (54). *Streptomyces* strains were grown at 30°C on DNA (2.3% Difco^TM^ nutrient agar), MH agar (1.5% agar in Mueller-Hinton broth, Oxoid) or MS agar (2% Mannitol, 2% Soya flour, 2% Agar) and in YEME broth (0.3% yeast extract, 0.3% malt extract, 0.5% peptone, 1% glucose, 34% sucrose). *E. coli* was cultivated in LB medium (1% Trypton, 0.5% Yeast Extract, 0.5% NaCl). For antibiotic selection, media were supplemented with the indicated antibiotics at the following concentrations: apramycin 50 mg/l, kanamycin 50 mg/l, carbenicillin 100 mg/l, chloramphenicol 25 mg/l, nalidixic acid 25 mg/l, and hygromycin at 40 mg mg/l for *Streptomyces*strains or 80 mg/l for *E. coli*.

For sporulating strains standard spore inoculum adjusted to a concentration of 0.5–0.6 McFarland was prepared by diluting spore suspensions in sterile 0,9% NaCl. For non-sporulating strains, overnight mycelium grown on agar plates was resuspended in sterile 0,9% NaCl. Minimum inhibitory concentrations (MICs) were determined by spotting 5 µl of standard inoculum onto MH agar plates containing a serial two-fold dilution range of antibiotics. MICs were evaluated after incubation at 30°C for 5 days. MIC testing was performed in at least three biological replicates for each strain. For co-cultivation experiments, *C. passeckerianus* CCBAS738 and CCBAS739 were grown on tryptic soy agar (TSA, Oxoid) at 28°C for 20 days. Agar discs approximately 8 mm in diameter were cut from the grown mycelium and transferred onto plates with the lawn of standard spore inoculum of *S. coelicolor* strain. Growth was evaluated after incubation at 30°C for 5 days.

For testing the GUS activity on agar plates, standard spore inoculum was streaked onto MS agar containing 160 mg/l of 5-bromo-4-chloro-3-indolyl-β-D-glucoronide (Sigma Aldrich)eand cultivated at 30 °C for the indicated time.

### Spectrophotometric GUS assays

For spectrophotometric GUS assays (55) the pellet from cultivation in YEME was resuspended in 1 ml of lysis buffer (50 mM phosphate buffer at pH 7, 0.1% triton X-100, 0.27% β-mercaptoethanol, and 4 g/l lysozyme), the mixture was incubated for 30 min at 37 °C and cell debris was removed by centrifugation. 25 μl of cell lysate was assayed in triplicate in a 96-well assay plate, mixed with 75 μl of phosphate buffer (8 g/l Na_2_HPO_4_.7H_2_O, 2.35 g/l NaH_2_PO_4_ H_2_O, pH 7) and 10 μl of 4 g/l p-nitrophenyl-β-D-glucuronide (Sigma Aldrich) added to each well. OD_420_ was measured at 30°C on a Tecan plate reader, the amount of total protein released from the mycelium was measured using Bradford reagent (BioRad).

### Cultivation for proteomic and metabolomic analysis

Spores of the tested strains were germinated for 8 h in 50 ml of 2xYT medium (54), each strain in triplicate. Germinated spores were centrifuged, and the pellet was resuspended in 5 ml of YEME medium, briefly sonicated (5s at 50% power on ice, Sonicator Hielscher UP220S), and divided into triplicates for inoculation into 50 ml of YEME medium to achieve a final spore concentration of OD_450_ 0.25–0.35. Cultivations were performed in 250 ml Erlenmayer flasks with metal springs. Upon visual detection of pink-colored prodiginines which appears in late exponential phase (at 18 h for Δ*tiaA* and Δ*are5sc* mutants, and 22 h for the WT strain), cultures were induced with the specified antibiotics. Samples (1 ml) were collected at 24 h and 84 h or 108 h, centrifuged to separate the pellet and supernatant, and stored at -80°C for subsequent proteomic (pellet) and metabolomic (supernatant) analyses.

### Proteomic analysis

#### Protein Digestion

Cell pellets were resuspended in 50 µl of 100 mM TEAB (Triethylammonium bicarbonate) containing 2% SDC (sodium deoxycholate) and boiled for 5 min. Protein concentration was determined using BCA protein assay kit (Thermo) and 30 µg of protein per sample was used for MS sample preparation. To reach total volume of 40 µl 100 mM TEAB containing 2% SDC, 40mM chloroacetamide, 10mM TCEP (Tris(2-carboxyethyl)phosphine) was added and heated for 5 min at 95°C. Samples were further processed using SP3 beads according to (56). Briefly, 5 µl of SP3 beads was added and filled to 50 µl with 100mM TEAB. Protein binding was induced by addition of ethanol to 60 % (vol./vol.) final concentration. Samples were mixed and incubated for 5 min at RT. After binding, the tubes were placed into magnetic rack and the unbound supernatant was discarded. Beads were subsequently washed two times with 180 µl of 80% ethanol. After washing, samples were digested with trypsin (trypsin/protein ratio 1/30) reconstituted in 100mM TEAB at 37°C overnight. After digestion samples were acidified with TFA to 1% final concentration and peptides were desalted using in-house made stage tips packed with C18 disks (Empore) according to (57).

### nLC-MS 2 Analysis

Nano Reversed phase columns (EASY-Spray column, 50 cm x 75 µm ID, PepMap C18, 2 µm particles, 100 Å pore size) were used for LC/MS analysis. Mobile phase buffer A was composed of water and 0.1% formic acid. Mobile phase B was composed of acetonitrile and 0.1% formic acid. Samples were loaded onto the trap column (C18 PepMap100, 5 μm particle size, 300 μm x 5 mm, Thermo Scientific) for 4 min at 18 μl/min loading buffer composed of water, 2% acetonitrile and 0.1% trifluoroacetic acid. Peptides were eluted with Mobile phase B gradient from 4% to 35% B in 60 min. Eluting peptide cations were converted to gas-phase ions by electrospray ionization and analyzed on a Thermo Orbitrap Fusion (Q-OT-qIT, Thermo Scientific). Survey scans of peptide precursors from 350 to 1400 m/z were performed in orbitrap at 120K resolution (at 200 m/z) with a 5 × 10^5^ ion count target. Tandem MS was performed by isolation at 1.5 Th with the quadrupole, HCD fragmentation with normalized collision energy of 30, and rapid scan MS analysis in the ion trap. The MS2 ion count target was set to 10^4^ and the max injection time was 35 ms. Only those precursors with charge state 2–6 were sampled for MS2. The dynamic exclusion duration was set to 45 s with a 10 ppm tolerance around the selected precursor and its isotopes. Monoisotopic precursor selection was turned on. The instrument was run in top speed mode with 2 s cycles (58).

### Proteomic data analysis

All data were analyzed and quantified with the MaxQuant software (version 2.1.0.0) (59). The false discovery rate (FDR) was set to 1% for both proteins and peptides and we specified a minimum peptide length of eight amino acids. The Andromeda search engine was used for the MS/MS spectra search against the S*treptomyces coelicolor* Reference Uniprot proteome (UP000001973, containing 8 038 entries). Enzyme specificity was set as C-terminal to Arg and Lys, also allowing cleavage at proline bonds and a maximum of two missed cleavages. Carbamidomethylation of cysteine was selected as fixed modification and N-terminal protein acetylation and methionine oxidation as variable modifications. The “match between runs” feature of MaxQuant was used to transfer identifications to other LC-MS/MS runs based on their masses and retention time (maximum deviation 0.7 min) and this was also used in quantification experiments. Quantifications were performed with the label-free algorithm in MaxQuant (60). Data analysis was conducted on biological triplicates using Perseus 1.6.15.0 software (61). Significantly altered protein levels between antibiotic-treated and untreated samples of the same strain were identified using a paired Student’s t-test, with a permutation-based approach to control the False Discovery Rate (FDR). Protein annotations from UniProt were complemented with KEGG annotations and custom annotations for known biosynthetic gene clusters proteins (62), SigR regulon (26) and WblC regulon (15). Enrichment analysis of these protein annotations was performed separately for upregulated and downregulated proteins using Fisher’s exact test, with Benjamini-Hochberg FDR correction applied (FDR ≤ 0.02).

The mass spectrometry proteomics data has been deposited at PXD059630.

### Metabolomic analysis

#### Solid phase extraction

Analyzed metabolites were extracted from the culture supernatants using solid phase extraction that was carried out as follows. An Oasis MCX 3cc 60 mg cartridge (Waters, USA) was conditioned with 3 mL methanol, equilibrated with 3 mL 2% formic acid (formic acid, 98– 100%, Honeywell, USA) and then 3 mL culture supernatant (pH adjusted to 3 with, formic acid, 98– 100%) was loaded. Thereafter, the cartridge was washed with 3 mL 2% formic acid and finally eluted with 1.5 mL methanol, followed by a wash with 2 mL methanol. Finally, a second elution with 1.5ml methanol with 1.45% ammonium hydroxide (Honeywell, USA). Both the eluents were evaporated to dryness (Concentrator Plus, 2013 model, Eppendorf), reconstituted in 300 μL 50% methanol and centrifuged at 13,000 × g for 5 min.

#### Liquid-Liquid extraction

Liquid-Liquid extraction was performed using ethyl acetate (Lach:ner, 99.89%) according to the protocol reported by (28). An equal volume of culture broth and ethyl acetate was added in a vial and vortexed for 10 s. The vials were then sonicated for 10 min and agitated at 200 rpm for 10 min at 28 °C, followed by centrifugation at 5000 rpm for 10 min at 4 °C. The layer with ethyl acetate was collected and filtered through the 0.2 µm polypropylene VWR® syringe filters. Then, ethyl acetate was evaporated with speed vacuum (Concentrator Plus, 2013 model, Eppendorf), and the sample was reconstituted in DMSO:methanol (1:9 v/v) LC-MS grade, and centrifuged at 13,000 × g for 5 min. It was done mainly to detect actinorhodin and undecylprodigiosin. To detect undecylprodigiosin the samples were further diluted 10 times using MeOH and 5 µL was used for injection in UPLC-Q-TOF-MS.

#### UPLC-Q-TOF-MS, UPLC-UV analysis and data processing

LC-MS analysis was performed on a Waters Acquity M-class UHPLC system connected to a Synapt G2-Si Q-TOF mass spectrometer (Waters Corporation, Manchester, UK). The LC column used was ACQUITY UPLC® BEH C18 (1.0 mm × 100 mm) 1.7µm column. 5 µL of the sample was injected into the column, which was kept at 40 °C. A two component mobile phase, B and A, acetonitrile (J.T.Baker LCMS grade) and 0.1% formic acid (Serva LC-MS grade, 99%) in water, respectively, were used for the elution at the flow rate of 80 µL/min. The elution was performed with condition as (min/%B) 0/5; 1.5/5; 15/70; 19/100 followed by 3min column clean-up (100% B) and 3 min equilibration (5% B). The mass spectrometer operated in the positive mode with capillary voltage +3kV, desolvation gas temperature, 250°C, cone voltage +40V; Source temperature, 70°C; cone gas flow, 50 L h^−1^; desolvation gas flow, 600 L h^−1^; scan time of 0.15 s; survey inter-scan time of 0.01 s. The lock spray technology was used to maintain the mass accuracy. 0.01% sodium trifluoroacetate was used for mass calibration, and fenclonine spiked in the samples at 3 ug/mL was used as internal standard. The mass spectra were collected in the 50–1700 m/z range with mass accuracy below 1ppm. The samples were injected in a randomized order. QC samples prepared as a mixture of all extracts at equal volumes were injected after every 6 injections.

The data processing was performed using Waters MassLynx V4.2 (Manchester, UK). The raw data files were converted to .mzML format and then it was used to process in a program to perform scan sequential and mass correction (using lockmass 556.2771). The final .mzML files were used for further processing. The visualization of the data was performed using SeeMS from Proteowizard (63). It was used to visualize the MS/MS fragments of each pseudomolecular ion corresponding to the detected metabolites. Finally, the MS/MS fragmentation spectra obtained experimentally were compared with the GNPS Library (http://gnps.ucsd.edu), Mona (https://mona.fiehnlab.ucdavis.edu/) and CFM-ID spectral prediction (https://doi.org/10.1021/acs.analchem.1c01465) in the case of unavailability of the fragmentation spectra in the aforementioned spectral libraries, in order to support the metabolite identity beyond accurate mass and fitting isotopic pattern of these metabolites that are not commercially available as standards. Analyzed metabolites as listed in Supplementary Table 6 (64–66). Moreover, the area of the peak was calculated for each pseudomolecular ion [M+H] using integration option available (Peak-to-peak amplitude 2000, with enable smoothing option on) in MassLynx and it was documented.

For Actinorhodin, the column used was the Acquity Premier CSH C18 column (2.1 mm × 50 mm I.D., particle size 1.7 μm, Waters). 5 µl of the sample was injected, which was kept at 40 °C. Two component mobile phase, B and A, acetonitrile (J.T.Baker LC-MS grade) and 0.1 % formic acid in water (Serva LCMS grade, 99%), respectively, were used for the elution at the flow rate of 0.4 ml/min. The elution was performed with condition as (min/% B) 0/5; 1.5/5; 10/46; 15/100 followed by 1 minute clean-up with 100% B and 3 minutes equilibrium with 5% B. LC-MS analyses were performed on Acquity UPLC system with 2996 Photo Diode Array detector. Absorption spectra of the actinorhodin was recorded from 190 to 700 nm. For actinorhodin we used standards with different concentrations (1000 µg to 7.625 µ g/ml) to obtain a calibration curve. The peak of the actinorhodin was detected at 520 nm. The calibration curve was used to perform absolute quantification and to obtain the area under the peak. The data processing was done using MassLynx V4.1 software.

Lastly, the area of the peak for all the analyzed metabolites were used for statistical analysis using Perseus software version 2.0.10.0 (https://maxquant.org/perseus). To identify metabolites significantly altered by antibiotic treatment, area of the peak values was Log2-transformed and Z-score normalized, missing values were imputed by zero and paired Student’s t-test, with a permutation-based approach to control FDR was performed. All the acquired LC-MS files, and metadata tables along with predicted spectra from CFM ID were deposited as .xls, .raw, .pdf and .mzML files (https://doi.org/10.5281/zenodo.14643682; https://doi.org/10.5281/zenodo.14644045) in the Zenodo (https://zenodo.org/).

### Toeprinting assay

Toeprinting was performed as described previously (67) with minor modifications. The detailed protocol can be found in the Supplementary Methods. All DNA templates (Supplementary Table 7) for *in vitro* transcription contained T7 promoter and the NV1 sequence (68). Template *are5sc* 5’UTR_L1 coding for *are5sc*L1 uORF (MLV peptide) with its natural ribosome binding site combined with artificial downstream sequence in order to prevent unwanted secondary structures of synthetized mRNA. Template *are5sc* uORF_L1 (AUG_mut_) differed by mutation changing start codon from TTG to ATG. Template are5sc 5’UTR contained *are5sc* 5’UTR_L1 with start codon mutated to ATG, and native downstream sequence including also predicted *are5sc* 5’UTR_L2 coding for the LRS peptide. Template *tiaA* 5’UTR contained native 5’UTR sequence coding for *tiaAL* (MVGDDDISG peptide). *In vitro* transcribed mRNA template at a final concentration of 1 μM was translated *in vitro* in the presence of antibiotics in 5 μl final reaction volume using PURExpress Δ Ribosome Δ Release Factors Kit (New England Biolabs). Ribosomes were purified from *Streptomyces venezuelae* ATCC 1071 as for this organism standardized methods for in vitro translation are available (Toh et al. 2021b2021), (Supplementary Methods). The tested antibiotics lincomycin, clindamycin, pristinamycin IIA, tiamulin, or erythromycin were dried in the tubes using a vacuum concentrator (Eppendorf) prior to the translation reaction in the amount corresponding to a final concentration in the reaction.

### Construction of ARE5 phylogenetic tree

ARE5 protein sequences were extracted from the existing database (9). The database was enriched with ARE5 protein sequences from the tested *Actinomycetota* strains. For strains that had not been sequenced, the ARE5 sequence was retrieved from another strain of the same species. Protein sequences were aligned using MAFFT version v7.526 with FFT-NS-i strategy. For removing poorly aligned regions (with more than 60% gaps) TrimAl (v1.4.rev15) was used with -gt option set on 0.4. After concatenation of alignment, strains with identical sequences or sequences that contain more than 45% of gaps were removed from the database. The resulting database contains 401 sequences with 538 positions. The phylogenetic tree was reconstructed using RAxML-NG (v.1.2.2). Specifically, we used LG+I+G4 substitution model, favoured by ModelTest-NG (v.0.1.7), with 200 bootstrap replicates (‘-seed 568317’ option). The phylogenetic tree was visualized using iTOL (v.6.9).

## Acknowledgments

This research was supported by the Ministry of Education, Youth and Sports of the Czech Republic through the grant “Talking Microbes – Understanding Microbial Interactions within the One Health Framework” (CZ.02.01.01/00/22_008/0004597), by the project National Institute of Virology and Bacteriology (Programme EXCELES, ID Project No. LX22NPO5103), funded by the European Union1Next Generation EU, as well as the OP JAK - MSCA Fellowships CZ (CZ.02.01.01/00/22_010/0006118). The research project was also supported by the Grant Agency of Charles University – grant number 134324 (granted to M.N.) and 1767418 (granted to L.V.). Proteomic analyses were conducted in the Laboratory of Mass Spectrometry at the BIOCEV Research Center, Faculty of Science, Charles University. We kindly acknowledge RNDr. Miroslav Flieger from the Institute of Microbiology, ASCR, for the isolation of actinorhodin, and RNDr. Ivana Eichlerová, Ph.D., from the Culture Collection of Basidiomycetes of the Institute of Microbiology, ASCR, for providing the *C. passeckerianus*strains CCBAS738 and CCBAS739. We also acknowledge to Dr. Simon J. Moore for providing the plasmid p1486 and the strain *S. venezuelae* ATCC 10712. Additionally, we thank Bc. Jan Jareš for facilitating the computational requirements necessary for the phylogenetic analyses.

## Author Contributions

M.K. and L.V. constructed most of the strains, performed most of the experiments

M.N. and D. M. did the toeprinting assay

A.M. and Z.K. performed the metabolomic analysis

N.P. Participated in preparation of mutant strains

J.P. and P.O.P. participated in GUS assay analysis

G.B.N. designed and supervised the experiments, G.B.N. and M.K. wrote the manuscript with consultation with other coauthors.

Z.K. revised the text

