## Supplemental material for "ABCF Protein-Mediated Resistance Shapes Bacterial Responses to antibiotics Based on their Type and Concentration"

[#] authors contributed to the manuscript equally

*****corresponding author

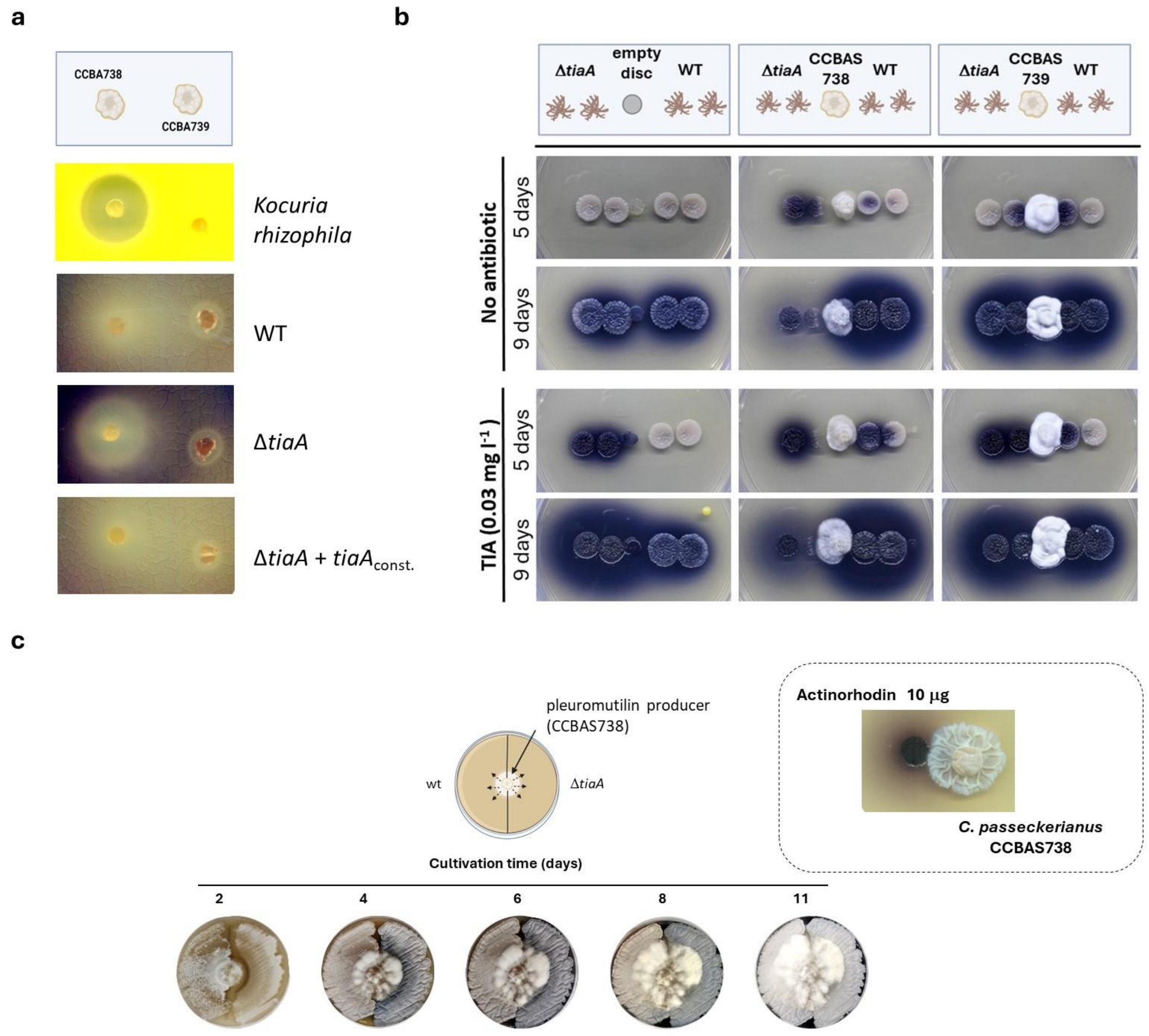


**Supplementary Figure 1. *Streptomyces coelicolor* and fungi *Clitopilus passeckerianus* co-cultivation experiments show that TiaA affects susceptibility and actinorhodin production in response to pleuromutilin produced by fungi (a)** *C. passeckerianus* strain CCBAS738 that produce pleuromutilin inhibits growth of pleuromutilin susceptible *Kocuria rhizophila* and *tiaA* mutant of *S. coelicolor*. The sensitive phenotype of *tiaA* mutant is complemented by the constitutive expression of *tiaA*. Inhibition of *S. coelicolor* growth by *C. passeckerianus* CCBAS739, which does not produce pleuromutilin is not dependent on the presence of *tiaA*. **(b)** Actinorhodin (actinorhodin) production in the presence of *C. passeckerianus* strains that produce (CCBAS738) and do not produce (CCBAS739) pleuromutilin on MH agar. The subinhibitory concentration of pleuromutilin supplemented in the agar induces earlier production of actinorhodin in the absence of fungi. Pleuromutilin produced by CCBAS738 induces earlier ($5^{th}$ day) but weak production of actinorhodin in *tiaA* mutant. Actinorhodin production does not differ between WT and *tiaA* mutant when co-cultured with CCBAS739. **(c)** Co-culture of WT and *tiaA* mutant with CCBAS738 on MS agar shows that production actinorhodin in *tiaA* mutant does not affect the growth of fungus which is also demonstrated by cultivating CCBAS738 next to the disk impregnated with 10 µg of purified actinorhodin.

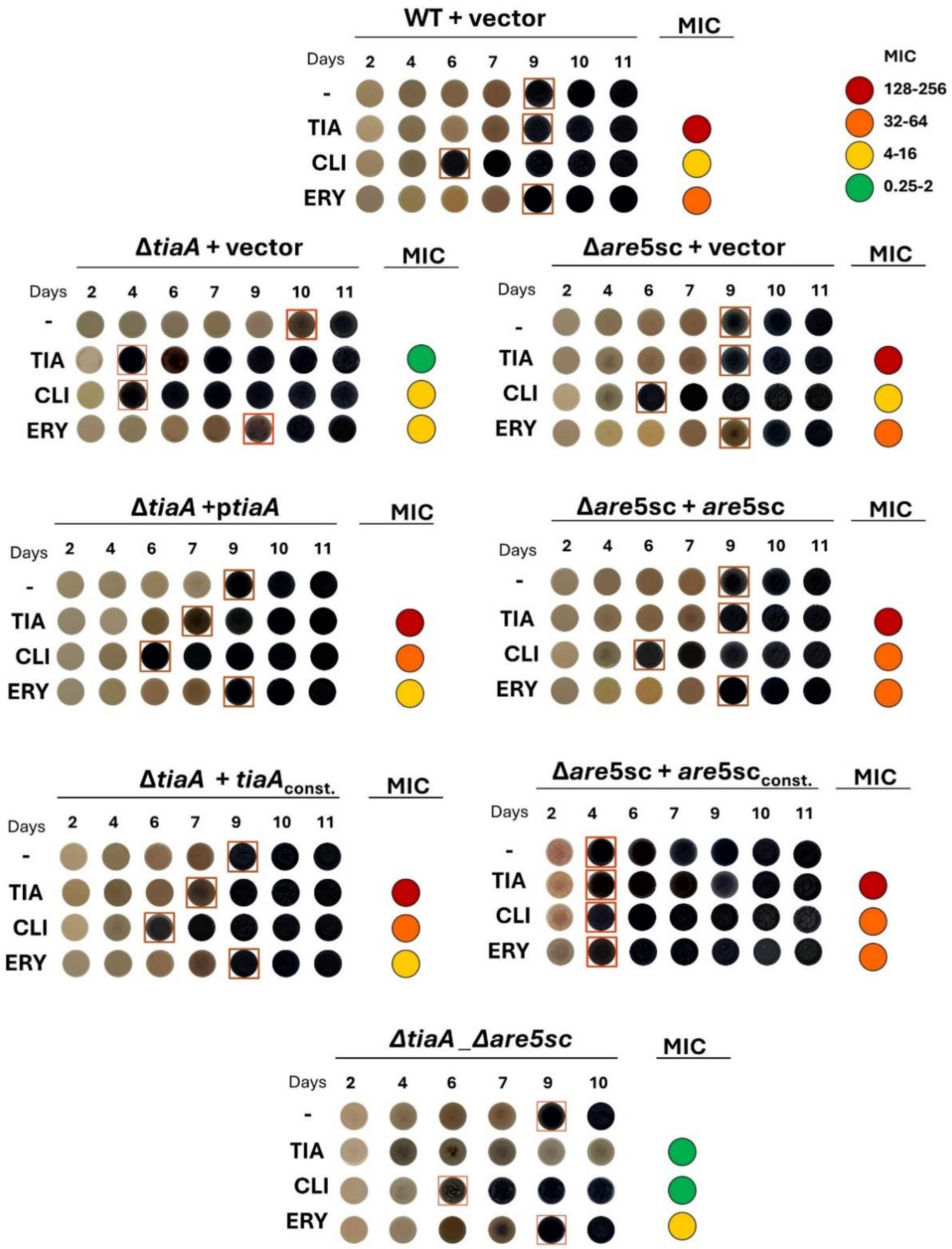


**Supplementary Figure 2. Impact of TiaA and Are5sc on actinorhodin production in response to antibiotics** S*treptomyces coelicolor* WT and mutant strains were cultured on MH agar supplemented with tiamulin (TIA), clindamycin (CLI) or erythromycin (ERY) at subinhibitory concentration 0.03 mg/l. Strains resistant to tiamulin due to *tiaA* expression exhibit delayed actinorhodin production. Overexpression of *are5sc* facilitates actinorhodin production independently of the antibiotic. The double mutant lacking both *tiaA* and *are5sc* fails to produce actinorhodin in response to antibiotics. Color coded MIC values are taken from Fig. 1a.

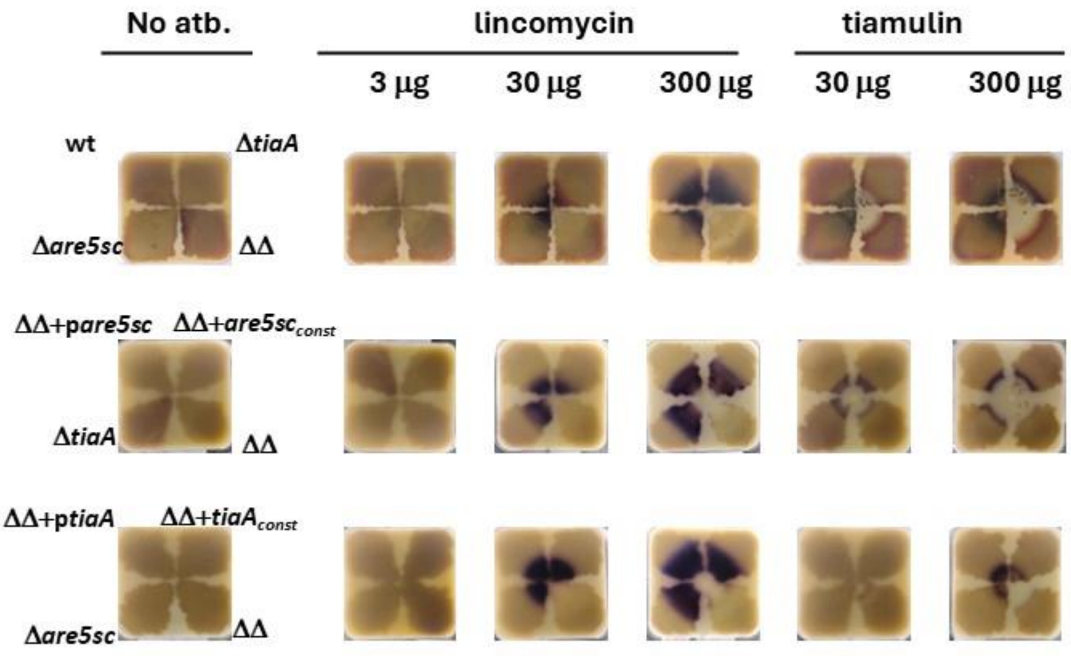


**Supplementary Figure 3. Impact of TiaA and Are5sc on actinorhodin production in response to lincomycin and tiamulin**. Antibiotic discs with varying concentrations were placed on MS agar plates to create concentration gradients. The experiment was repeated four times.

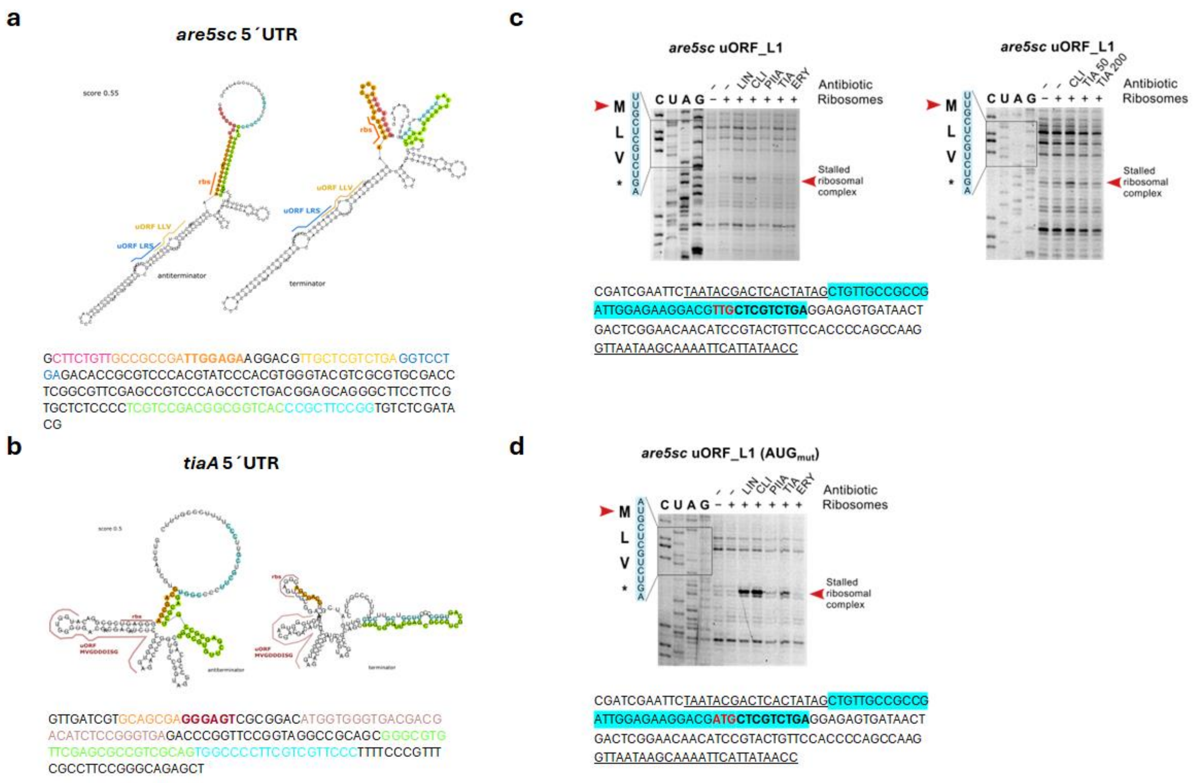


**Supplementary Figure 4.** Terminator and anti-terminator conformations of **(a)** are5sc and **(b)** tiaA 5´UTRs as predicted by PASIFIC(1). **(c)** Toeprinting assay performed on non-mutated *are5scL1* showing stalled ribosome complex (SRC) formed only by lincomycin (LIN, 50 µM) and clindamycin (CLI, 5 µM) but not by tiamulin (TIA, 50 µM) and pristinamycin IIA (PIIA, 30 µM) (left). Even an increase of TIA (200 µM) did not stall ribosomes (right). The template sequence corresponding to 5´UTR region including native ribosome binding site and are5scL1 is highlighted **(d)** Changing the start codon of *are5scL1* from ttg to *atg* resulted in ribosome stalling also by TIA and PIIA. The discrepancy between the toeprinting results and the induction profiles of the reporter constructs may arise from differences in translation initiation efficiency at non-canonical start codons between *E. coli* and *Streptomyces* (2). The *ttg* start codon of *are5scL1* is among the strongest in the TX/TL translation system of *S. venezuelae* but is considered only "moderate" in *E. coli*, likely due to variations in codon usage and the corresponding tRNA pools between these organisms (3, 4). In the toeprinting experiments, we utilized ribosomes from *S. venezuelae* in combination with *E. coli*-derived components with native *E. coli* tRNA pool, which may have reduced the efficiency of the *ttg* codon in this hybrid system. This could explain the inability to capture the antibiotic-ribosome complex for certain antibiotics.

a

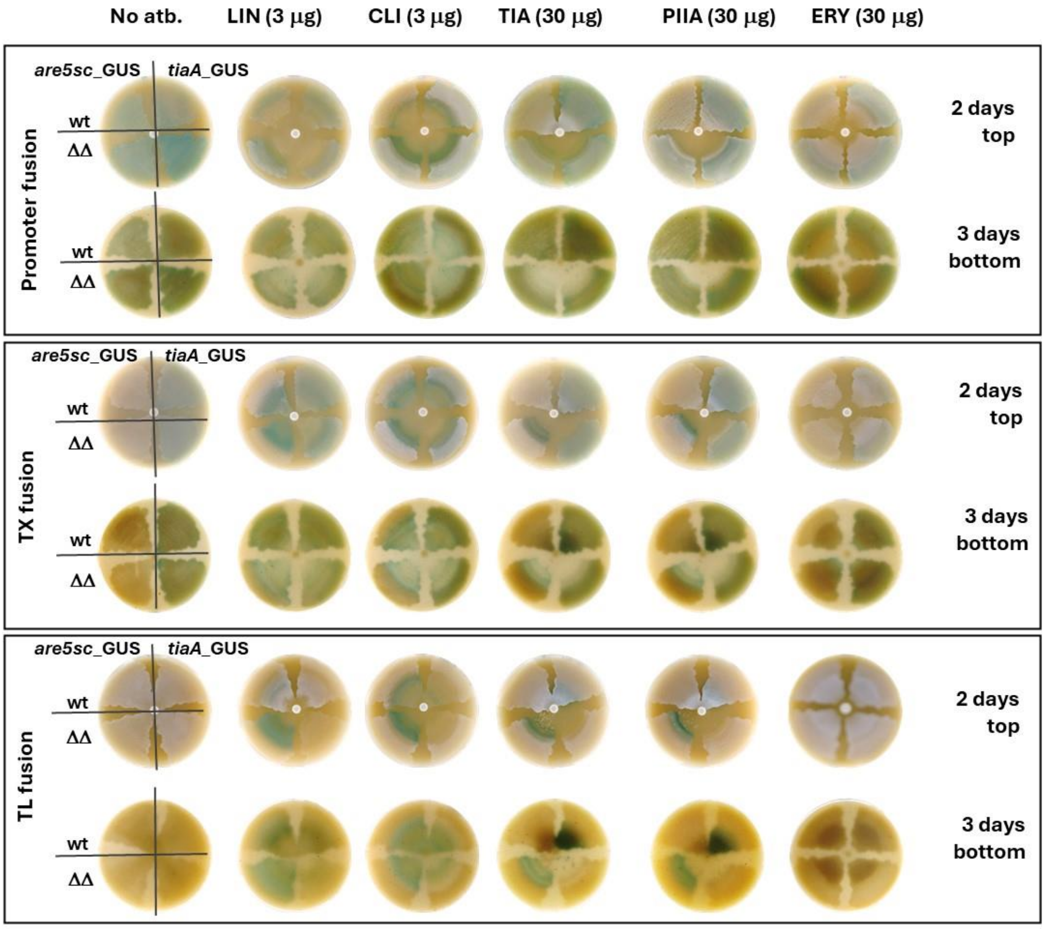


b

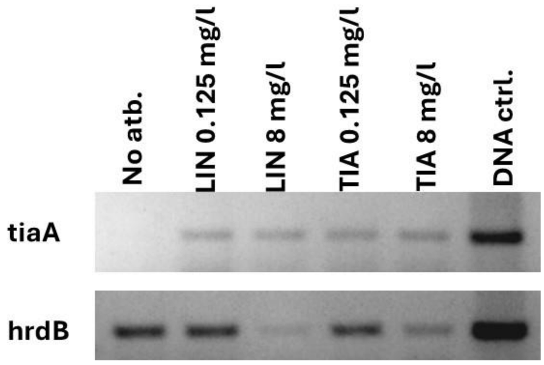


**Supplementary Figure 5. Expression of ARE5 proteins is regulated in response to antibiotics by transcriptional attenuation mechanism a)** Activities of β-glucuronidase (GUS) promoter (P), transcriptional (TX), and translational (TL) reporter fusions were monitored on plates with MS agar. The antibiotic diffusing from the disc in the center of the plate creates a concentration gradient. **P fusion reporters** show constitutive activity of *tiaA* and *are5sc* promoters, although antibiotic treatment initially inhibited the translation of the reporter gene, delaying GUS production. **TX fusion reporter** constructs showed that transcription of *are5sc* was induced by antibiotics, while *tiaA* was constitutive which is not consistent with the predicted transcriptional attenuation mechanism. Previous RNA-seq data sets indicated low-level constitutive transcription of *tiaA* that was induced by lincomycin (3, 5). It suggests that *tiaA* 5' UTR has a leaky premature terminator which in combination with a weak endogenous ribosome binding site, would result in low-level basal expression in the absence of antibiotics in the natural system. However, we used a strong synthetic ribosome binding site in our transcriptional fusions (6) that probably overrides this natural regulation, leading to constitutive activity of the reporter. Nevertheless, the increased GUS production around tiamulin and pristinamycin IIA discs in strains with TX-reporters indicate the transcriptional activation of TiaA expression **b)** RT-PCR analysis of *tiaA* transcript showing that *tiaA* is induced by antibiotics at transcriptional level. The *hrdB*, encoding the principal sigma factor was used as a control. For the analysis cultures of *S. coelicolor* WT were induced with antibiotics during the late exponential phase (40 h of cultivation), and the mycelium for RNA isolation was harvested 3 hours post-induction. Total RNA was isolated and reverse-transcribed into cDNA. PCR was then performed on the cDNA using specific primers tiaA for and tiaA rev to test the presence of *tiaA* transcript and hrdBF2 and hrdBR2 to test the presence of control gene *hrdB (*primers are listed in Supplementary table 5).

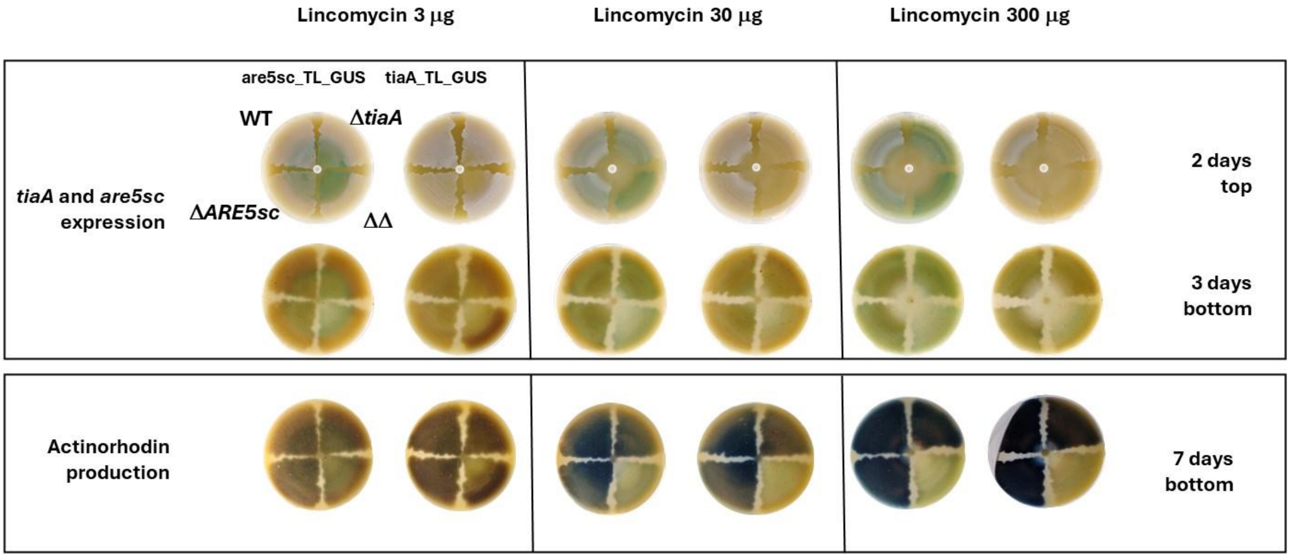


**Supplementary Figure 6. The presence of ARE5 proteins is essential for the induction of actinorhodin triggered by lincomycin.** The induction of actinorhodin production by lincomycin is concentration-dependent and requires the presence of ARE5 proteins. In the double mutant, this induction does not occur.

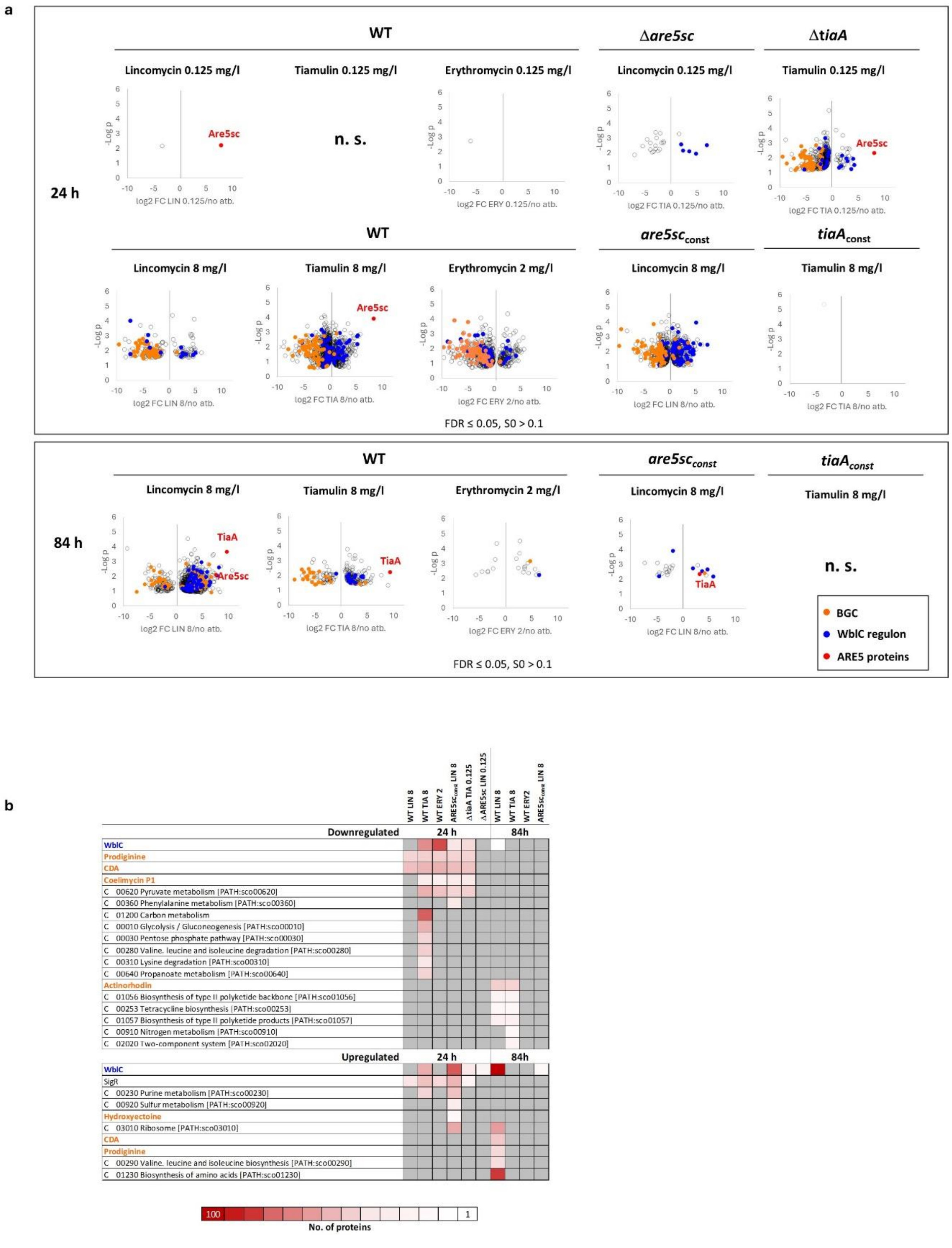


**Supplementary Figure 7. Proteomics analysis of *S. coelicolor* strains treated with antibiotics. (a)** Volcano plot showing significantly altered proteins in response to antibiotic treatment (FDR ≤ 0.05, S0 > 0.1) in low and high concentrations in WT, *tiaA,* and *are5sc* mutants and overexpression strains. Proteins involved in specialized metabolite biosynthesis (orange) and WblC (blue) regulon are highlighted **(b)** Results of Kyoto Encyclopedia of Genes and Genomes (KEGG), WblC (7) and SigR (8) regulons enrichment analysis of significantly downregulated and upregulated proteins in response to antibiotic treatment (FDR ≤ 0.05, S0 > 0.1) using a Fisher's exact test with a p-value threshold of 0.05 performed in Perseus (version 1.6.15). The heatmap shows the number of enriched proteins (intersection size) results of the full analysis are in the Source file).

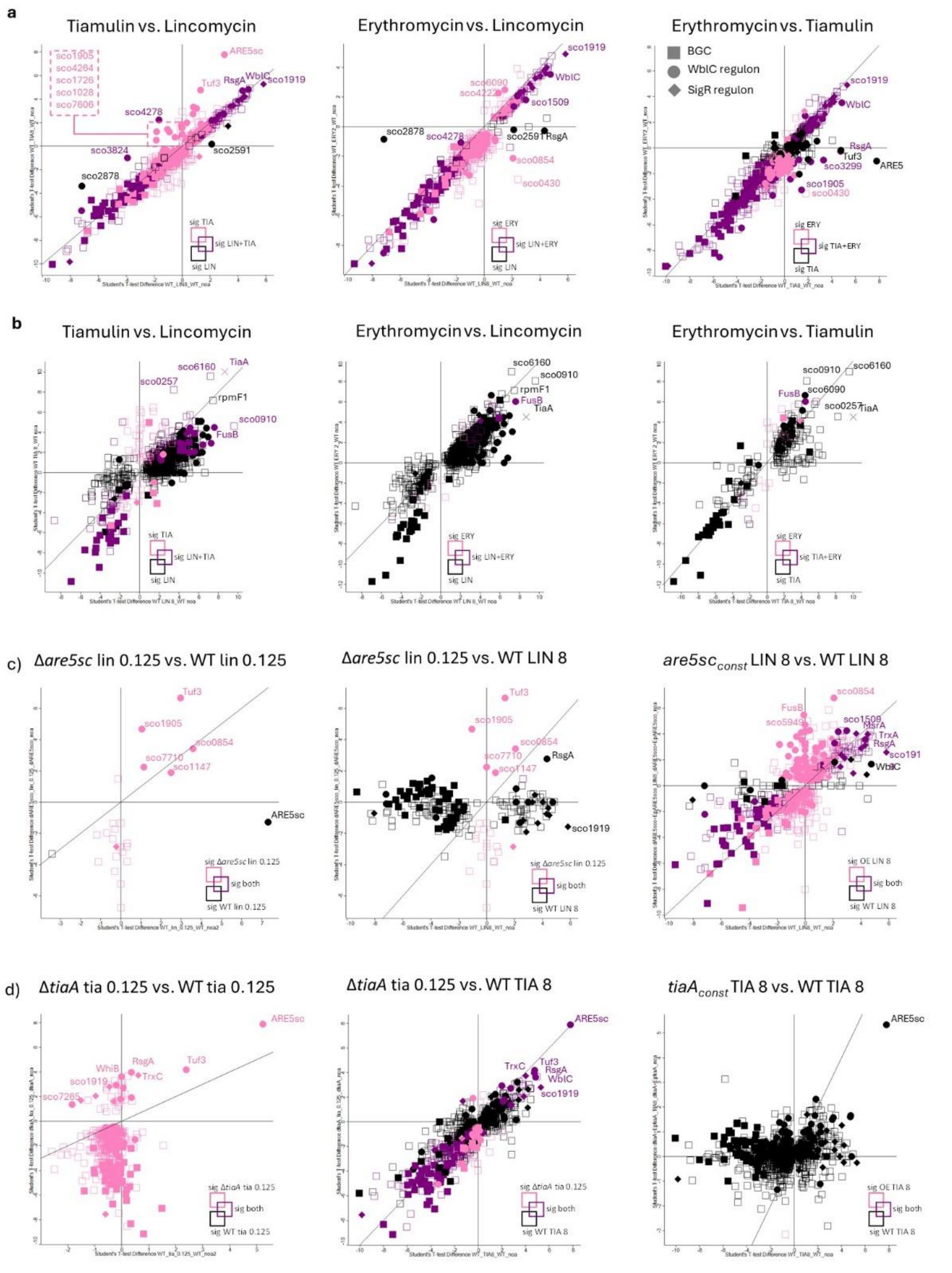


**Supplementary Figure 8. Comparison of changes in protein abundance in response to antibiotic treatment** Scatterplots comparing the fold change in protein abundance between antibiotic-treated and untreated samples. The x-axis shows the fold change in one analysis and the y-axis shows the fold change in the other analysis. Proteins significantly changed in one (pink or black) or both (violet) analyses are color-coded as indicated. **(a)** Comparison of wild-type response to tiamulin, lincomycin, and erythromycin at 24 hours **(b)** Comparison of wild-type response to tiamulin, lincomycin, and erythromycin at 84 hours **(c)** Comparison of wild-type and *are5sc* mutant/overexpression strains treated with lincomycin at 24 hours **(d)** Comparison of wild-type and *tiaA* mutant/overexpression strains treated with tiamulin at 24 hours **(e)** Comparison of wild-type and overexpression strains in 84 hours.

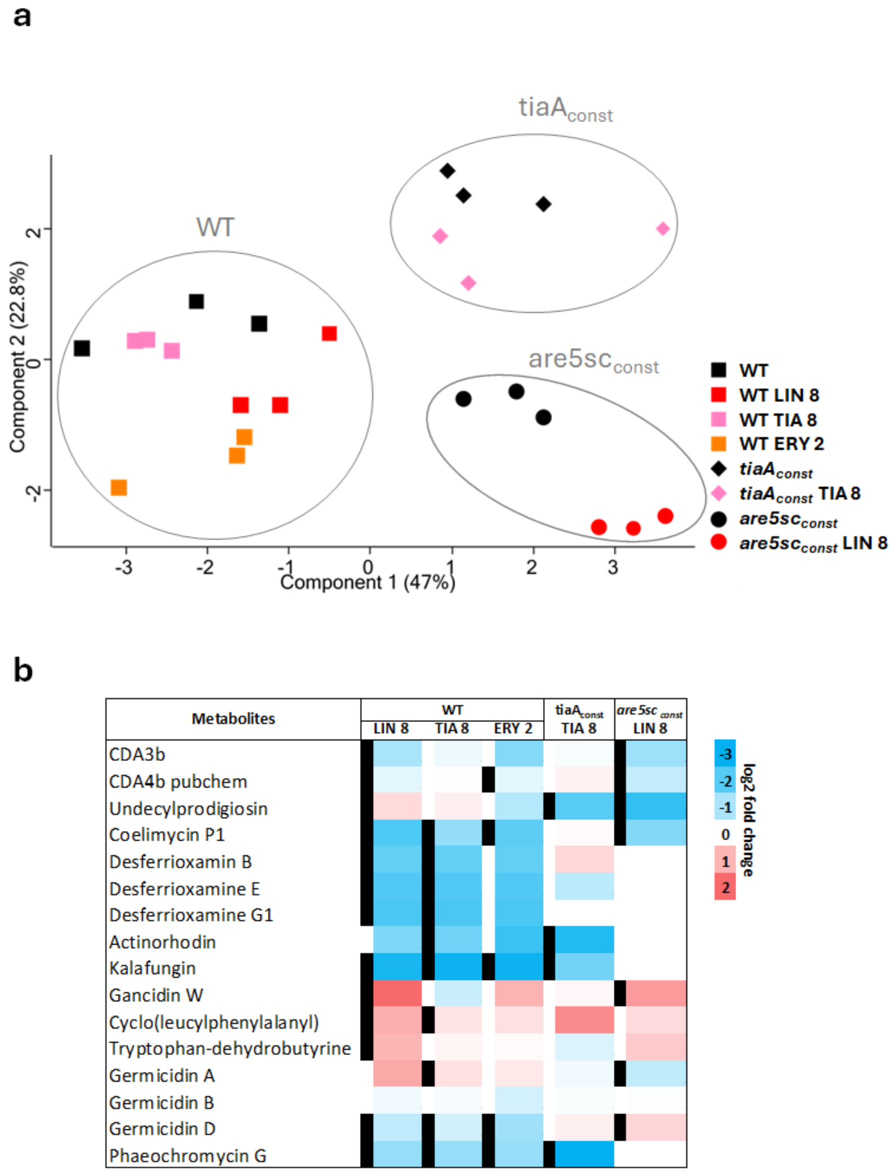


**Supplementary Figure 9. Antibiotic-induced changes in specialized metabolite production (a)** Principal component analysis of relative metabolite abundance, based on normalized peak intensities, detected in supernatants of wild-type and overexpression strains treated with high concentrations of antibiotics. (b) Significant changes in the abundance of known *S. coelicolor* secondary metabolites in response to antibiotic treatment. The relative abundance of metabolites corresponds to normalized peak intensities and were analyzed using paired Student's t-test with a false discovery rate (FDR) of ≤ 0.05 and S0 > 0.1.

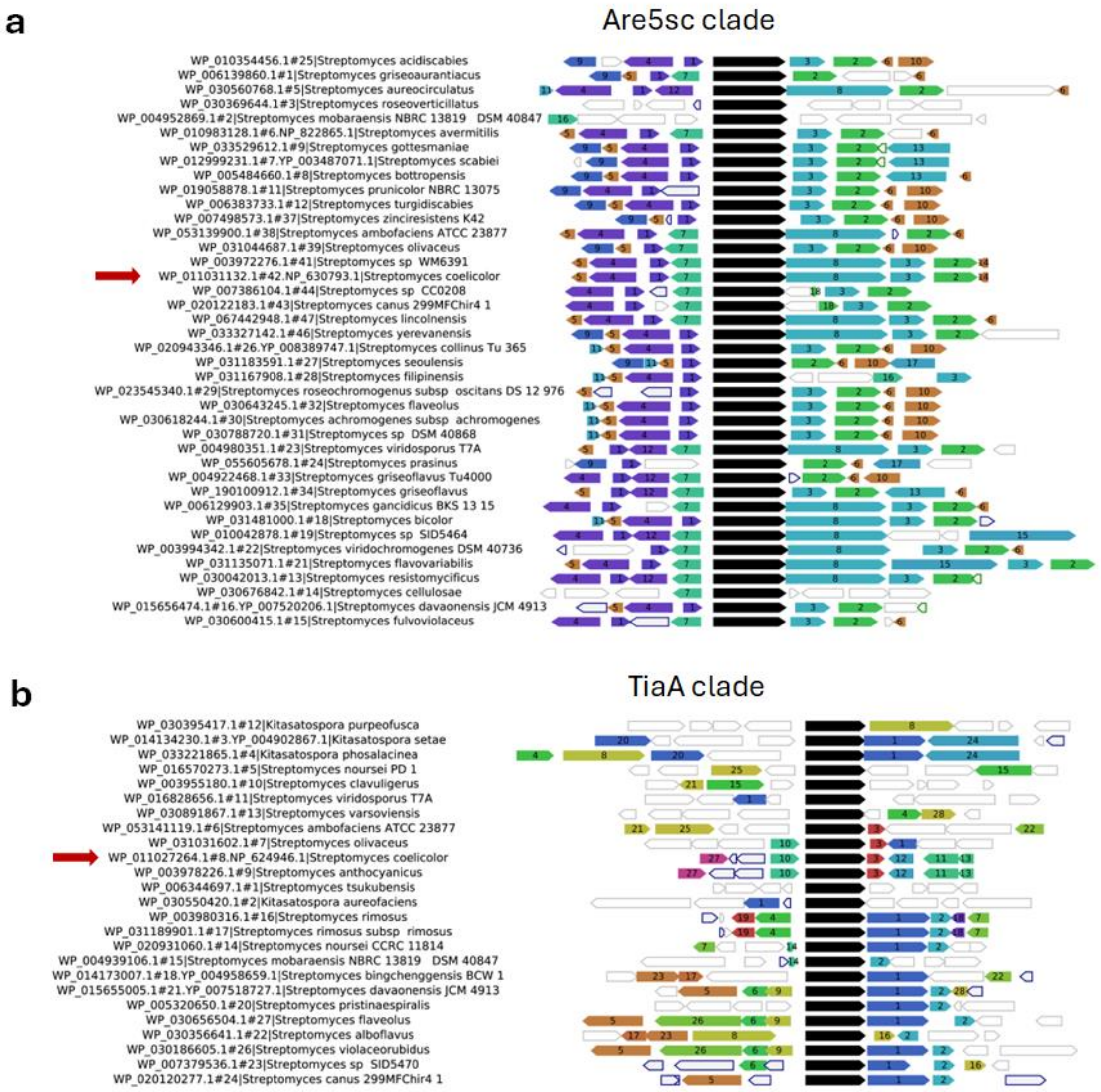


**Supplementary Figure 10. Genome context of Are5sc homologues is more conserved than context TiaA homologues** Results of the WebFlaGs (9)analysis shows genomic neighbourhood surrounding the gene encoding the ARE5 proteins from the **(a)** from the clade Are5sc and **(b)** clade TiaA. Genes encoding homologous proteins are labelled by the same number and color. Red arrows show *S. coelicolor*

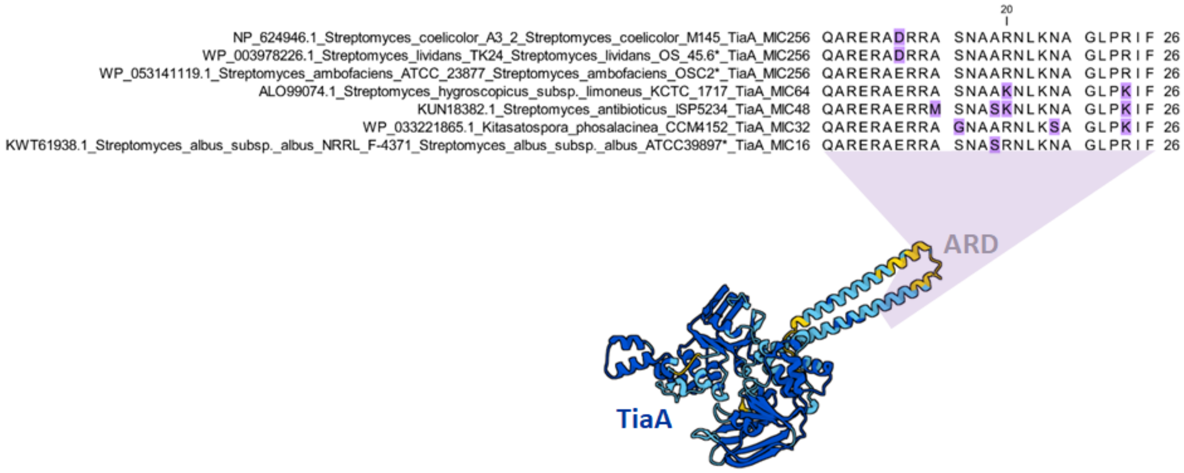


**Supplementary Figure 11. Alignment of amino acid sequences of TiaA homologues from *Actinomycetota* strains tested for tiamulin susceptibility.** The alignment shows that amino acid substitutions in TiaA homologs correlate with lower MIC values (16-64 mg/L). The aligned region corresponds to an extension of the antibiotic resistance domain (ARD), specifically at the end of the linker (also known as the PtMI domain) that interacts with the ribosomal antibiotic binding site. The region that was compared is indicated in the AlphaFold model (10, 11) of TiaA.

**Supplementary Table 1.** Susceptibility of mycelia of *S. coelicolor* M145 (WT), ARE5 deficient strains to lincomycin (LIN), clindamycin (CLI), tiamulin (TIA).

| | **$MIC_{MYCELIUM}$ (mg/l)** | | |
|---|---|---|---|
| | LIN | CLI | TIA |
| *S. coelicolor* + vector | 32 | 8 | 128 |
| *S. coelicolor* Δ*tiaA* + vector | 8 | 4 | 0.25 |
| *S. coelicolor* Δ*are5sc* + vector | 32 | 8 | 128 |

**Supplementary Table 2.** Susceptibility of spores of *Streptomyces lincolnensis ΔlmrC* constitutively expressing *tiaA* or *are5sc* to lincomycin (LIN), clindamycin (CLI), tiamulin (TIA), pristinamycin IIA (PIIA), erythromycin (ERY), chloramphenicol (CAM) and tetracycline (TET). The antibiotics are grouped based on resistance phenotypes conferred by ARE ABCF proteins: $LS_AP$ (lincosamides, streptogramin A, pleuromutilins), MSb (macrolides, streptogramin B), and PhO (phenicols, oxazolidinones).

| | $MIC_{SPORES}$ (mg/l) | | | | | | |
|---|---|---|---|---|---|---|---|
| | 50S | | | | | | 30S |
| | $LS_AP$ | | | | $MS_B$ | PhO | |
| | LIN | CLI | TIA | PIIA | ERY | CAM | TET |
| *S. lincolnensis ΔlmrC* + vector | >8192 | 256-512 | 2 | >64 | 32-64 | nd | nd |
| *S. lincolnensis ΔlmrC* + *tiaA* $_{const}$ | >8192 | 256-512 | 16 | >64 | 32 | nd | nd |
| *S. lincolnensis ΔlmrC* + *are5sc* $_{const}$ | >8192 | 128-256 | 0.5-1 | >64 | 16-32 | nd | nd |

### Supplementary Table 3. List of strains used in the study.

| Strain | Genotype/characteristic/use | Ref. |
|---|---|---|
| ***Escherichia coli*** **XL1** | Δ(*mcrA*)*183* Δ(*mcrCB-hsdSMR-mrr*)*173 endA1 supE44 thi-1 recA1* gyrA96 relA1 lac *[F'* proAB lac$I^{q}$ZΔ M15 *Tn*10 *(Tet$^{r}$)* | Stratagene |
| ***Escherichia coli*** **ET12567 pUZ8002** | *dam, dcm, hsdS, cat, tet*;carries plasmid pUZ8002 used for conjugative transfer between *E.coli* and *Streptomyces* spp. | Gust, B., Challis, G. L., Fowler, K., Kieser, T. & Chater, K. F. PCR-targeted Streptomyces gene replacement identifies a protein domain needed for biosynthesis of the sesquiterpene soil odor geosmin. Proc. Natl. Acad. Sci. U. S. A. 100, 1541–6 (2003). |
| ***Escherichia coli*** **BW25113 pIJ790** | $Cm^{r}$; K12 derivative (*lacI$^{q}$ rrnB*$_{T14}$ *ΔlacZ*$_{WJ16}$ *hsdR514 ΔaraBAD*$_{AH33}$ *ΔrhaBAD*$_{LD78}$) containing temperature sensitive vector pIJ790 for λ RED mediated recombination | Gust, B. *et al.* λ Red-mediated genetic manipulation of antibiotic-producing Streptomyces. *Adv. Appl. Microbiol.* **54**, 107–128 (2004). |
| ***Escherichia coli*** **DH5α/BT340** | $Cm^{r}$, $Carb^{r}$; K12 derivate containing temperature sensitive vector for FLP recombination | Gust, B. *et al.* λ Red-mediated genetic manipulation of antibiotic-producing Streptomyces. *Adv. Appl. Microbiol.* **54**, 107–128 (2004). |
| **WT** | *Streptomyces coelicolor* M145, type strain, $SCP1^{-}$ $SCP2^{-}$ | Bentley, S. D. et al. Complete genome sequence of the model actinomycete Streptomyces coelicolor A3(2). Nature 3, 141–147 (2002). |
| **ΔtiaA** | *Streptomyces coelicolor M145 Δsco0636::apra* | This study |
| **Δare5sc** | *Streptomyces coelicolor M145 Δsco6720* | This study |
| **ΔtiaA_Δare5sc** | *Streptomyces coelicolor M145 Δsco0636::apra; Δsco6720* | This study |
| ***Streptomyces lincolnensis*** **ATCC 25466** | | |
| ***Streptomyces lividans*** **OS 45.6** | | |
| ***Streptomyces ambofaciens*** **OSC2** | | |
| ***Streptomyces hygroscopicus*** **subsp.** ***limoneus*** **KCTC 1717** | | |
| ***Streptomyces antibioticus*** **ISP 5234** | | |
| ***Kitasatospora phosalacinea*** **CCM 4152** | | |
| ***Streptomyces albus*** **subsp.** ***albus*** **ATCC 39897** | | |
| ***Streptomyces globisporus*** **DSM 40991** | | |
| ***Streptomyces griseus*** **CCM 3178** | | |
| ***Streptomyces niveus*** **DSM 40088** | | |
| ***Streptomyces catenulae*** **DSM 40258** | | |
| ***Nocardiopsis prasina*** **DSM 43845** | | |
| ***Streptomyces griseoflavus*** **Tu4000** | | |
| ***Streptomyces griseoflavus*** **NRRL B-5312** | | |
| ***Streptomyces prasinus*** **NRRL B-12521** | | |
| ***Streptomyces caelestis*** **ATCC 15084** | | |
| ***Streptomyces avermitilis*** **DSM 46492** | | |
| ***Amycolatopsis keratiniphila*** **subsp.** ***keratiniphila*** **NRRL B-24117** | | |
| ***Nocardia carnea*** **CCM 2756** | | |
| ***Nocardia tenerifensis*** **DSM 44704** | | |
| ***Streptomyces albus*** **J1074** | | |
| ***Streptomyces scabrisporus*** **DSM 41855** | | |
| ***Micrococcus luteus*** **DSM 20030** | | |
| ***Kocuria rhizophila*** **CCM 552** | | |
| ***Streptomyces venezuleae*** **ATCC 10712** | | Moore, Simon J., et al. "A Streptomyces venezuelae cell-free toolkit for synthetic biology." ACS synthetic biology 10.2 (2021): 402-411. |
| ***Clitopillus passeckerianus*** **CCBAS 738** | strain producing pleuromutilin | Culture Collection of Basidiomycetes (CCBAS) https://www.biomed.cas.cz/ccbas/fungi.htm |
| ***Clitopillus passeckerianus*** **CCBAS 739** | | Culture Collection of Basidiomycetes (CCBAS) https://www.biomed.cas.cz/ccbas/fungi.htm |

### Supplementary Table 4. List of oligonucleotides used in the study.

| Oligonucleotide | Sequence |
|---|---|
| sgRNA_R | ACGCCTACGTAAAAAAAGCACCGACTCGGTGCC |
| sgRNA_6720_F | CATGCCATGG ATGCCCAGCGACTCCAGGAAGTTTTAGAGCTAGAAATAGC |
| Stu1(5)_6720_F | TCGTCGAAGGCACTAGAAGGGAGCCCGATCGCCTCCACCC |
| Stu1(3)_6720_R | GGTCGATCCCCGCATATAGGGTGACGGACGCCGCTCAGCC |
| HR_6720_F | CTCATCCAGCGGACGCTAGGCGGAATACCCGGGGGAAGTAGACATAGGGGTCTCCGCGGT |
| HR_6720_R | ACCGCGGAGACCCCTATGTCTACTTCCCCCGGGTATTCCGCCTAGCGTCCGCTGGATGAG |
| Sco0636pIJFW | TCCTCATATGTCCGACGCCGCTGTC |
| Sco0636pIJRev | ATATCTCGAGCTACTCCTCCGGCGCC |
| Sco6720pIJFW | AACAAGCTTCTAGGCGGAATACCCG |
| Sco6720pIJRev | CCTTCATATGTCTACTTCCCCCACTTCTCTC |
| pG109 | GCCGCCGATTGGAGAAGGACGTTGCTCGTCAATGAACCGTTGGAGGCAAACACATATG |
| pG094 | AAACAGCTATGACATGATTACGAATTCGATTCACTGCTTGCCGCCCTG |
| pG110 | GACGAGCAACGTCCTTCTCCAATC |
| pG096 | ATCGAATTCGTAATCATGTCATAGCTGTTTCCTGTG |
| pG111 | GCCGGGAAGTCCACCCTGTTGAAGCTGATCtgaAATGAACCGTTGGAGGCAAACACATATG |
| pG094 | AAACAGCTATGACATGATTACGAATTCGATTCACTGCTTGCCGCCCTG |
| pG112 | GATCAGCTTCAACAGGGTGGACTTC |
| pG120 | CCGGGCGGCGGCTCCATGCTGCGCCCGGTCGAAAC |
| pMK034 | ATGGTGACGAAGGAACTACTAGTTAGC |
| pMK035 | GCTAACTAGTAGTTCCTTCGTCACCATTCACTGCTTGCCGCCCTG |
| pG105 | GTCGCGGACATGGTGGGTGACGACGACATCAATGAACCGTTGGAGGCAAACACATATG |
| pG106 | GATGTCGTCGTCACCCACCATG |
| pG107 | GCCGGGAAGTCCACCCTGCTCAAGCTGATCtgaAATGAACCGTTGGAGGCAAACACATATG |
| pG108 | GATCAGCTTGAGCAGGGTGGACTTC |
| pMK13 | AGCGGGCAGGGAGCGGATCCGCGGCCGCGCGCGATCCACCTGCTCGCACCGGAAG |
| pMK15 | ACAGGAAACAGCTATGACATGATTACGAATTCGATTGTGATGCACGATCAGCAAAG |
| pMK012 | AGCGGGCAGGGAGCGGATCCGCGGCCGCGCGCGATCAGTGAGAGCGAGCCGTTG |
| pG053 | ACAGGAAACAGCTATGACATGATTACGAATTCGATCTAGGCGGAATACCCGACAG |
| pG085 | AGATCTCCAGGATCCCCTGGATAATTAATTAATGAACCGTTGGAGGCAAACACATATGCTGCGCCCGGTCGAAAC |
| pG087 | TCCGCTCATGAGAACCTAGGATCCAAGCTTTCACTGCTTGCCGCCCTG |
| pG086 | AATTAATTATCCAGGGGATCCTGGAGATC |
| pG084 | TGAAAGCTTGGATCCTAGGTTCTCATG |
| FP_T7_TP | CGATCGAATTCTAATACGACTCACTATAG |
| RP_NV1_TP | GGTTATAATGAATTTTGCTT |
| NV_YY | GGTTATAATGAATTTTGCTTATTAAC |
| tiaA for | CTGATGGACTACACCGGCTG |
| tiaA rev | CTGCGGACGTTCTTCTCGG |
| HrdB F2 | CGCGGCGTCGTCTCCATGC |
| HrdB R2 | TGCGAGCGCGAGGGGTGA |

### **Supplementary Table 5. List of plasmids used in the study.**(12–16)

| plasmid | Genotype/characteristic/use | Ref. |
|---|---|---|
| **pIJ773** | Apra$^r$ Carb$^r$; vector carrying inactivation cassette containing *aac(3)IV* gene of apramycin resistance and *oriT* surrounded by FRT region | Gust, B. et al. λ Red-mediated genetic manipulation of antibiotic-producing Streptomyces. Adv. Appl. Microbiol. 54, 107–128 (2004). |
| **pMS81** | Hyg$^r$; ϕBT1 *attP* integrative vector for conjugative transfer from *E.coli* to *Streptomyces* spp. | Gregory, M. a & Smith, M. C. M. Integration Site for Streptomyces Phage jBT1 and Development of Site-Specific Integrating Vectors. J. Bacteriol. 185, 5320–5323 (2003). |
| **pIJ10257** | Hyg$^r$; Integrative and conjugative vector derived from pMS81 with constitutive promoter *ermEp*, ribosome binding and multicloning sites | Hong, H. J., Hutchings, M. I., Hill, L. M. & Buttner, M. J. The role of the novel fem protein VanK in vancomycin resistance in Streptomyces coelicolor. J. Biol. Chem. 280, 13055–13061 (2005). |
| **cosmid 7G01** | SuperCos bearing *sco0636* and surrounding genes | https://strepdb.streptomyces.org.uk |
| **pCrisprCas9** | pGM1190-sgRNA with cas9 | Tong, Yaojun, et al. "CRISPR-Cas9 toolkit for actinomycete genome editing." Synthetic Metabolic Pathways: Methods and Protocols 163-184 (2018) |
| **p1486** | pTU1-A-SP44 plasmid expressing β-glucuronidase | Toh, Ming, et al. "A High-Yield Streptomyces TX-TL Toolkit for Synthetic Biology and Natural Product Applications." *Journal of visualized experiments: JoVE* 175 (2021): 10-3791. |
| **pGBN064** | Hyg$^r$ , pIJ10257 bearing region starting with of lmrC to the end of lmbU translationally fused with mCherry. | Koberska, Marketa, et al. "Beyond self-resistance: ABCF ATPase LmrC is a signal-transducing component of an antibiotic-driven signaling cascade accelerating the onset of lincomycin biosynthesis." MBio 12.5 (2021): 10-1128. |
| **pMK048** | pCrisprCas9 with sgRNA for *are5sc* knock out and two regions homologus to regions flanking *are5sc* . Region 1 was amplified from genomic DNA of *S. coelicolor* M145 using primers Stu1(5)_6720_F and HR_6720_R and region 2 was amplified using primers HR_6720_R and Stu1(3)_6720_R). Amplified regions were assembled using NEBuilder® HiFi DNA Assembly kit (NEBuilder, New England Biolabs) with pCrisprCas9 vector linearised by StuI. sgRNA was inserted into this plasmid using two step Golden Gate assembly cloning: sgRNA was amplified from pCrisprCas9asusing sgRNA_R and sgRNA_6720R, cut with BsaI and ligated in pCrisprCas9 digested with NcoI and NheI (as described in Tong, Yaojun, et al. "CRISPR–Cas9, CRISPRi and CRISPR-BEST-mediated genetic manipulation in streptomycetes." Nature Protocols (2020): 1-33.) | This study |
| **$tiaA_{const}$ (pLV032)** | Hyg$^r$ , pIJ10257 vector containing *tiaA* under constitutive *ermE* p* promoter was used for complementations. The plasmid was constructed by ligating a fragment amplified from the genomic DNA of *S. coelicolor* M145 using primers Sco0636pIJFW and Sco0636pIJRev into the vector pIJ10257. The fragment and vector were digested with the restriction endonucleases NdeI and XhoI prior to ligation. | This study |
| **ptiaA (pMK095)** | Hyg$^r$ , pMS81 containing gene sco0636 with its 400 bp upstream region used for complementations. The plasmid was constructed using the NEBuilder by combining EcoRV-linearized pMS81 with a fragment amplified from the genomic DNA of *S. coelicolor* M145 using primers pMK13 and pMK15. | This study |
| **$are5sc_{const}$ (pLV040)** | Hyg$^r$ , pIJ10257 containing gene sco6720 under ermEp promoter. Plasmid was prepared via ligation of the fragment (amplified on gDNA of *S. coelicolor* using primers Sco6720pIJFW and Sco6720pIJRev into vector pIJ10257 cut by NdeI and HindIII restriction endonucleases. | This study |
| **pare5sc (pMK090)** | Hyg$^r$ , pMS81 vector containing *are5sc* with its 323 bp upstream region was used for complementations. The plasmid was constructed using the NEBuilder by combining EcoRV-linearized pMS81 with a fragment amplified from the genomic DNA of S. coelicolor M145 using primers pMK12 and pG053. | This study |
| **pGBN129** | Hyg$^r$, pIJ10257 vector containing the gus N2RBS (AATGAACCGTTGGAGGCAAACACAT) followed by the β-glucuronidase gene from the plasmid p1486. The plasmid was prepared by NEBuilder by combining the insert amplified from plasmid p1486 pG085 and pG087 and the pGBN064 plasmid PCR-linearized with primers pG086 and pG084. | This study |
| **tiaA_P_GUS (pGBN142)** | Hyg$^r$ , pMS81 vector which includes a 259 bp long region that contains the promoter of tiaA, followed by N2 ribosome-binding site (N2RBS; AATGAACCGTTGGAGGCAAACACAT) and the *gus* gene encoding β-glucuronidase. The plasmid was constructed using the NEBuilder by combining insert amplified from plasmid pGBN129 with primers pG105 and pMK035 and plasmid pMK095 linearized by PCR with primers pG106 and pMK034. | This study |
| **tiaA_TX_GUS (pGBN143)** | Hyg$^r$ , pMS81 vector containig the 400 bp upsteram region of *tiaA,* region encoding the first 51 aminoacids of TiaA and N2RBS and the *gus* . The plasmid was constructed using NEBuilder by combining the insert amplified from plasmid pGBN129 using primers pG107 and pMK035 and the pMK095 plasmid PCR-linearized with primers pG108 and pMK03. | This study |
| **tiaA_TL_GUS (pGBN144)** | Hyg$^r$ , pMS81 vector containig the 400 bp upsteram region of *tiaA* , region encoding the first 51 aminoacids of TiaA fused with the *gus* by the PGGGS linker (CCGGGCGGCGGCTCC). The plasmid was prepared by linearisation of the plasmid pGBN143 with using primers pG120 and pG108, subsequent phosphorylation of the 5' ends and self ligation. | This study |
| **are5sc_P_GUS (pGBN140)** | Hyg$^r$ , pMS81 vector containig the promoter region of *are5sc* , a N2RBS and the *gus* . The plasmid was constructed using NEBuilder by combining the insert amplified from plasmid pGBN129 using primers pG109 and pG094 and plasmid pMK090 PCR-linearised with primers pG110 and pG096 | This study |
| **are5sc_TX_GUS (pGBN141)** | Hyg$^r$ , pMS81 vector containig the 323 bp upsteram region of *are5sc* , region encoding the first 53 aminoacids of Are5sc and N2RBS and the *gus* by the PGGGS linker. The plasmid was constructed using NEBuilder by combining the insert amplified from plasmid pGBN129 using primers pG111 and pG094 and plasmid pMK090 PCR-linearized with primers pG112 and pG096. | This study |
| **are5sc_TL_GUS (pGBN145)** | Hyg$^r$ , pMS81 containig the 323 bp upsteram region of *are5sc* , region encoding the first 53 aminoacids of Are5sc fused with the *gus via* the PGGGS linker. The plasmid was prepared by linearisation of the plasmid pGBN141 with using primers pG120 and pG112, subsequent phosphorylation of the 5' ends and self ligation. | This study |

**Supplementary Table 6. List of compounds analyzed using LC-MS.**

| Name of the compound | M+H | MS/MS fragment match with GNPS library (*) or *in silico* prediction by CFM ID (#) or Mona (~) |
|---|---|---|
| Germicidin D | 169.0864 | 68.05, 79.05, 83.04, 91.05, 95.08, 105.07, 123.08, 153.05, 154.06, 169.08 (*) |
| Germicidin B | 183.1021 | 67.05, 79.05, 91.05, 94.08, 103.05, 107.05, 137.09, 153.05, 167.07, 168.08, 183.101(*) |
| Germicidin A | 197.1177 | 69.03, 77.04, 97.02, 105.06, 125.05, 135.04, 153.05, 167.07, 168.078, 197.117 (*) |
| gancidin W | 211.1446 | 58.02, 70.06, 72.04, 98.05, 138.13, 154.07, 211.14 (~) |
| phaeochromycin G | 217.0864 | 107.04, 115.05, 131.05, 133.02, 159.04, 173.06 (#) |
| cyclo(leucylphenylalanyl) | 261.1603 | 86.09, 91.05, 93.06, 103.05, 118.06, 120.08, 141.07, 231.11 (#) |
| tryptophandehydrobutyrine diketopiperazine (TDD) | 284.1399 | 103.05, 115.05, 117.06, 118.06, 130.06, 142.06, 144.08, 153.06, 253.09(#) |
| Kalafungin | 301.0712 | 85.03, 93.03, 121.03, 133.03, 149.02, 175.04, 177.01, 181.05, 187.03, 189.05, 201.06, 217.05 (#) |
| Coelimycin P1 | 349.1222 | 68.04, 117.05, 150.04, 176.05, 190.07, 204.05, 218.06 (*) |
| Desferrioxamin B | 561.3611 | 84.08, 102.09, 144.10, 165.10, 201.12, 202.12 (*) |
| Desferrioxamine E | 601.3561 | 84.08, 100.04, 120.07, 138.09, 165.10, 183.11, 201.12, 401.24 (*) |
| Desferrioxamine G1 | 619.3667 | 100.04, 154.08, 165.10, 184.10, 201.12, 401.24 (*) |
| CDA3b | 1483.529 | 55.05, 69.06, 85.06, 113.06, 118.06, 130.06, 132.08, 1281.43, 1298.46, 1310.42, 1311.45, 1338.46, 1354.45, 1371.47, 1391.51, 1413.49, 1423.51, 1437.52, 1447.56, 1465.52 (#) |
| CDA4b | 1497.5446 | 55.05, 60.04, 85.06, 118.06, 130.06, 132.08, 1298.46, 1310.46, 1322.46, 1325.46, 1340.47, 1354.49, 1385.49, 1406.52, 1427.50, 1433.53, 1451.54, 1461.52, 1478.53 (#) |
| undecylprodigiosin | 394.2858 | 159.05, 208.08, 224.08, 238.10, 240.11, 252.11, 280.15, 322.19, 350.22 (#) |

### Supplementary Table 7. Toeprint templates

| No. | Name | Coding sequence | Nucleotide sequence |
|---|---|---|---|
| **TPMN010** | ***are5sc* uORF_L1** | MLV | CGATCGAATTCTAATACGACTCACTATAGCTGTTGCCGCCGATTGGAGAAG<br>GACGTTGCTCGTCTGAGGAGAGTGATAACTGACTCGGAACAACATCCGTA<br>CTGTTCCACCCCAGCCAAGGTTAATAAGCAAAATTCATTATAACC |
| **TPMN022** | ***are5sc* uORF_L1 (AUGmut)** | MLV | CGATCGAATTCTAATACGACTCACTATAGCTGTTGCCGCCGATTGGAGAAG<br>GACGATGCTCGTCTGAGGAGAGTGATAACTGACTCGGAACAACATCCGTA<br>CTGTTCCACCCCAGCCAAGGTTAATAAGCAAAATTCATTATAACC |
| **TPMN023** | ***are5sc* 5'UTR (AUGmut)** | MLV, LRS | CGATCGAATTCTAATACGACTCACTATAGCTGTTGCCGCCGATTGGAGAAG<br>GACGATGCTCGTCTGAGGTCCTGAGACACCGCGTCCCACGTATCCCACG<br>TAAATACGTCGCGTGCGACCTGTTAATAAGCAAAATTCATTATAACC |
| **TPMN019** | ***tiaA* 5'UTR** | MVGDDDISG | CGATCGAATTCTAATACGACTCACTATAGCGTTGATCGTGCAGCGAGGGA<br>GTCGCGGACATGGTGGGTGACGACGACATCTCCGGGTGAGACCCGGTTC<br>CGGTAGGCCGCAGCGGGCGTGTTAATAAGCAAAATTCATTATAACC |

### SUPPLEMENTARY METHODS

#### Toeprinting assay

*Toeprinting samples preparation.* Commercially acquired DNA template was amplified by PCR using the primers FP_T7_TP (CGATCGAATTCTAATACGACTCACTATAG) and RP_NV1_TP (GGTTATAATGAATTTTGCTT). DNA template was transcribed *in vitro* using T7 RNA Polymerase (New England Biolabs) and NTPs (Thermo Fisher Scientific) according to the manufacturer's instructions and purified by RNA Clean & Concentrator Kit (Zymo Research). Either pure DMSO or antibiotics dissolved in DMSO were evaporated in 1.5 ml tube before the beginning of the translation reaction. mRNA template at a final concentration of 1 µM was translated *in vitro* in 5 µl reaction volume in the presence of lincomycin, clindamycin, pristinamycin IIA, tiamulin, erythromycin or tetracycline at different final concentrations using PURExpress Δ Ribosome Δ Release Factors Kit (New England Biolabs) and ribosomes at a final concentration of 2.4 µM purified from *S. venezuelae*. After incubating for 20 min at 37 °C, 1 µl of Yakima Yellow-5'-labelled probe NV_YY (2 µM) complementary to the NV1 sequence (GGTTATAATGAATTTTGCTTATTAAC) was added to the reaction and the sample was incubated for another 5 min at 37 °C. Reverse transcription was then performed with 5 U of Avian Myeloblastosis Virus reverse transcriptase (Promega Corporation) mixed with 0.4 µl of Pure System Buffer (9 mM Magnesium acetate, 5 mM Potassium phosphate, 95 mM Potassium glutamate, 5 mM $NH_4Cl$, 0.5 mM $CaCl_2$, 1 mM spermidine, 8 mM putrescine, 1 mM DTT) and 0.1 µl of dNTP mix (Thermo Fisher Scientific; each 10 mM) for 20 min at 37 °C. RNA was degraded by adding 0.5 µl of a 10 M NaOH stock at 37 °C for 15 min. Samples were neutralized with 0.7 µl of a 7.5 M HCl stock and 20 µl of Toeprinting Resuspension Buffer was added (0.3 M Sodium acetate, pH 5.5, 5 mM EDTA, pH 8, 0.5% SDS). cDNA was purified using a Nucleotide Removal Kit (QIAGEN) and eluted with 50 µl of water. After evaporation, fully dried samples were resuspended in 3 µl of formamide loading dye (95% formamide, 250 µM EDTA, pH 8, 0.25% Bromphenol blue).

*Sanger sequencing.* Sequencing reactions were performed with 1 pmol of DNA template treated with ExoSAP-IT reagent (Thermo Fisher Scientific) using Hemo KlenTaq polymerase (New England Biolabs), $MgCl_2$ at a final concentration of 0.6 mM and Yakima Yellow-5'-labelled probe NV_YY at a final concentration of 0.12 µM added to a tube containing 20 µM dNTP mix and one of the ddNTP (30 µM ddGTP, 350 µM ddATP, 600 µM ddTTP, 200 µM ddCTP, respectively). The following thermocycler program was used: 60 s of initial denaturation at 95 °C; 40 cycles of 30 s of denaturation at 95 °C, 30 s of annealing at 42 °C, and 60 s of elongation at 70 °C; and an additional 5 min of elongation at 70 °C. Sequencing reactions were purified by precipitation with 2% lithium chloride in acetone and resuspended in 3 µl of formamide loading dye.

*Gel analysis.* Sequencing reactions were denatured at 75 °C for 3 min, the toeprinting samples were denatured at 95 °C for 5 min and both were analysed on a 6% sequencing polyacrylamide gel (42 cm height, 20 width, 0.4 mm depth) in TBE buffer (10.8% Trizma Base, 5.5% boric acid, 20 mM EDTA, pH 8). The gel was run 2.5 h at 40 W and imagined on Typhoon™ FLA 9500 (GE Healthcare Life Sciences).

*Purification of ribosomes.* The 70S ribosomes were purified from 1 l of the *S. venezuelae* ATCC 10712 culture cultivated at 30°C (200 rpm) until reaching $OD_{600}$ value of 2.4. The cells were suspended in buffer A (100 mM Tris-HCl, pH 7.5, 500 mM $NH_4Cl$, 52.5 mM $MgCl_2$, 2.5 mM EDTA) supplemented with a protease cocktail inhibitor tablet (Sigma-Aldrich). Subsequently, cell lysis was achieved through five cycles of cryomilling (Retsch) 5 x 3 min cycles, speed 30 1/s, initial cooling 8 min, speed 5 1/s, intermittent cooling cycles 1 min. The resulting cell lysate was clarified by centrifugation at 38,000 x g at 4°C for 1 hour and then filtered (0.45 µm). The clarified cell lysate was layered onto a 37.7% sucrose cushion in buffer A and centrifuged at 100,000 × g for 16 h at 4°C using a fixed angle rotor 50.2-Ti for ribosome pelleting. The ribosome pellet was washed in buffer A, followed by centrifugation at 10000 × g for 5 minutes, and subsequently resuspended in buffer B (100 mM Tris-HCl, pH 7.5, 2500 mM $NH_4Cl$, 75 mM $MgCl_2$, 2.5 mM EDTA). The ribosomes were pelleted at 50,000 RPM for 2 hours at 4°C using a fixed angle rotor 50.2-Ti. This ribosome pellet was dissolved in a storage buffer (20 mM Tris-HCl, pH 7.5, 100 mM $NH_4Cl$, 12.5 mM $MgCl_2$, 0.5 mM EDTA, and 6 mM β-mercaptoethanol). The 70S ribosomes were fractionated on a 10-35% sucrose gradient equilibrated in buffer A and centrifuged at 38,694 × g for 16 hours using a Beckman SW41 Ti rotor. The resulting 70S ribosome peaks were collected *via* a piston fractionator (Biocomp). Subsequently, the 70S particles were washed with a storage buffer and then concentrated.
